# Targeting cellular senescence alleviates bone marrow aging

**DOI:** 10.64898/2026.07.16.736849

**Authors:** Bowen Yan, Jin Han, Yang Yang, Peiyi Zhang, Jason O. Brant, Jianhui Chang, Ha-Neui Kim, Maria Almeida, Prabhjot Kaur, Qingchen Yuan, Marco Demaria, Kalyanee Shirlekar, Jennifer Elisseeff, Guangrong Zheng, Ying Liang, Daohong Zhou, Olga A. Guryanova

## Abstract

Aging of the hematopoietic system impairs hematopoietic stem cell (HSC) function and alters bone marrow niche behavior, increasing susceptibility to anemia, infections, and hematologic malignancies. Here, pharmacologic clearance of senescent cells with the PROTAC compound 753b simultaneously targeting BCL-xL and BCL-2 reverses key secretory, transcriptional, and functional hallmarks of hematopoietic aging with low toxicity, restoring balanced lineage output. Single-cell RNA sequencing further demonstrates that 753b treatment attenuates aging-associated transcriptional signatures in HSCs, while selectively eliminating senescent, pro-survival niche cells without grossly perturbing niche composition. Functionally, 753b suppresses pro-inflammatory cues from both niche and hematopoietic cells including those emanating from neutrophil progenitors, rebalancing global bone marrow secretory ecosystem across stromal and hematopoietic compartments. Collectively, we identify 753b-induced senescent cell clearance as a powerful strategy to rejuvenate aged hematopoiesis and re-establish homeostatic communication between HSCs and their microenvironment, with implications for mitigating age-related hematologic dysfunction and improving hematologic health in older individuals.

## Main

Aging of the hematopoietic system compromises blood cell production, immune competence, and regenerative capacity, predisposing older individuals to anemia, infections, and hematologic malignancies^1–3^. Central to this decline is the functional deterioration of hematopoietic stem cells (HSCs), which progressively lose self-renewal potential, acquire myeloid bias and aberrant inflammatory signaling^2^. The bone marrow (BM) niche is crucial for HSC function and contributes to both HSC aging and rejuvenation^4,5^. Accumulating evidence indicates that persistent activation of stress and senescence pathways, due at least in part to pro-inflammatory BM microenvironment, disrupts HSC-niche homeostasis and drives age-associated alterations in hematopoiesis^6–10^.

Cellular senescence, a state of irreversible growth arrest marked by elevation of the cyclin-dependent kinase inhibitors p16INK4A (p16) and/or p21Cip1/Waf1 (p21), upregulation of anti-apoptotic proteins such as BCL-XL and BCL-2, and acquisition of the senescence-associated secretory phenotype (SASP), has emerged as a key mechanism driving tissue aging^3,11^. The accumulation of senescent cells with age impairs tissue repair and regeneration, promoting various age-related diseases through SASP-mediated chronic inflammation^12^. Proof-of-principle experiments in genetically engineered mouse models such as p16-3MR^13^ and *INK-ATTAC*^14^ that enable inducible ablation of senescent cells have demonstrated that depletion of senescent cells delayed onset of age-related functional deterioration within various tissues and extended health span^14,15^. Consistently, pharmacologic clearance of senescent cells, termed senolytic therapy, delays age-related symptoms and pathology in preclinical models^16–21^, demonstrating that pharmacologic senolysis is feasible in principle. However, first-generation senolytics have fallen short because of modest potency, suboptimal selectivity, therapy resistance, and dose-limiting hematologic toxicities, which hinder long-term treatment necessary for efficacy^21,22^.

The anti-apoptotic proteins BCL-2 and BCL-XL have been implicated in protecting senescent cells from pro-apoptotic signals and multiple BCL-2 family inhibitors have been developed^23–25^. Of these, venetoclax (a BCL-2 selective inhibitor) is the only FDA approved BCL-2 family inhibitor in the clinic but its use is still limited by rapid resistance development^26^, while navitoclax (a dual BCL-2/BCL-XL inhibitor) causes profound thrombocytopenia due to on-target BCL-XL inhibition in platelets, severely restricting its clinical utility^21,27,28^. To overcome these limitations, our team previously developed 753b, a novel BCL-2/BCL-XL proteolysis targeting chimera (PROTAC) that recruits both targets to the von Hippel-Lindau (VHL) E3 ligase for ubiquitylation and degradation^29,30^. Consequently, 753b spares platelets, which lack VHL expression and thus escape on target platelet toxicity characteristic of navitoclax^29,31^. These features provide a strong rationale to evaluate 753b as a highly effective next generation senolytic agent with improved hematologic safety profile in aged individuals.

Here, we comprehensively evaluate the efficacy of 753b, in restoring hematopoietic health in both aged HSCs and BM niche cells. By integrating *in vivo* functional assays with single-cell transcriptomic profiling of young, aged vehicle-treated, and aged 753b-treated mice, we demonstrate that 753b administration selectively clears senescent cell populations, resets aging-associated transcriptional programs in HSCs towards a more youthful state, restores balanced lineage output, and partially normalizes paracrine niche signaling between stromal and hematopoietic compartments. Our findings support 753b as a safe and effective senolytic suitable to rejuvenate hematopoiesis. More broadly, our study establishes pharmacologic senolysis as a powerful strategy to alleviate BM aging and to ameliorate age-associated hematopoietic stem cell dysfunction.

## Results

### Senescent cell clearance alleviates aging-associated functional decline of HSCs in a genetic p16-3MR mouse model

To investigate the impact of senescent cell clearance on HSC aging, we employed the p16-3MR transgenic mouse model, which enables selective elimination of p16-expressing senescent cells upon treatment with ganciclovir (GCV)^13^ (Fig. 1A). Twelve-months old mice were repeatedly treated with GCV or vehicle control until 24 months, at which point tissues were analyzed (Fig. 1B). Successful depletion of senescent cells was validated by quantitative RT-PCR analysis of adipose tissues to evaluate expression levels of senescence marker genes *Cdkn2a, Cdkn1a* and inflammatory cytokines *Il6*, *Il1b*, and *Mmp3* in young, aged vehicle-treated, and aged GCV-treated mice (Fig. 1C, SFig. 1A). As expected, we found significant elevation of these transcripts with aging, whereas with GCV treatment their expression was markedly reduced, confirming clearance of senescent cells and attenuation of tissue inflammation (Fig. 1C, SFig. 1A). Additionally, immunofluorescence staining of fluorescence-activated cell sorting (FACS)-sorted immunophenotypic HSCs revealed increased proportion of p16- or phospho-p38-positive HSCs in aged vehicle-treated mice, consistent with enhanced cellular senescence and stress signaling during aging in HSCs (Fig. 1D). In contrast, with GCV treatment the frequency of these senescent HSCs significantly decreased. Furthermore, immunofluorescence detection of γH2A.X foci similarly showed a significant reduction in DNA damage accumulation in HSCs in GCV-treated aged group compared to aged vehicle controls (Fig. 1D), suggesting successful senescent cell clearance. To further examine whether senescent cell depletion improves aged HSC function, we performed competitive bone marrow transplantation (BMT) assays (Fig. 1E). Fifty long-term HSCs (LT-HSCs) isolated from young, aged vehicle, or aged GCV-treated CD45.2-expressing donors were transplanted into lethally irradiated CD45.1-expressing recipients alongside 3×10^5^ unfractionated CD45.1^+^ bone marrow (BM) cells. Four months post transplantation, donor-derived hematopoietic reconstitution was assessed in peripheral blood by CD45.1/CD45.2 chimerism. Recipients of aged vehicle-treated donor HSCs exhibited significantly reduced donor cell chimerism compared to young controls, whereas aged GCV-treated donor HSCs showed substantially improved BM reconstitution (Fig. 1F, SFig. 1B). Analysis of mature lymphoid and myeloid lineages revealed marked myeloid skewing and reduced abundance of T and B lymphocytes in recipients of aged vehicle donor cells, characteristic of aged HSC dysfunction. In contrast, GCV treatment restored more balanced lymphoid-to-myeloid lineage output, comparable to young HSC donors (Fig. 1G, SFig. 1B). To directly test functional HSC frequency, we performed limiting dilution transplantation assays using purified LT-HSCs from each group (SFig. 1C). Four months post transplantation, we found a dramatic reduction in repopulation-competent HSC frequency in aged vehicle-treated donors (1/83) compared to young donors (1/9), whereas senescent cell clearance with GCV significantly increased frequency of functional HSCs in aged mice (1/11) (SFig. 1D, E).

**Figure 1.**
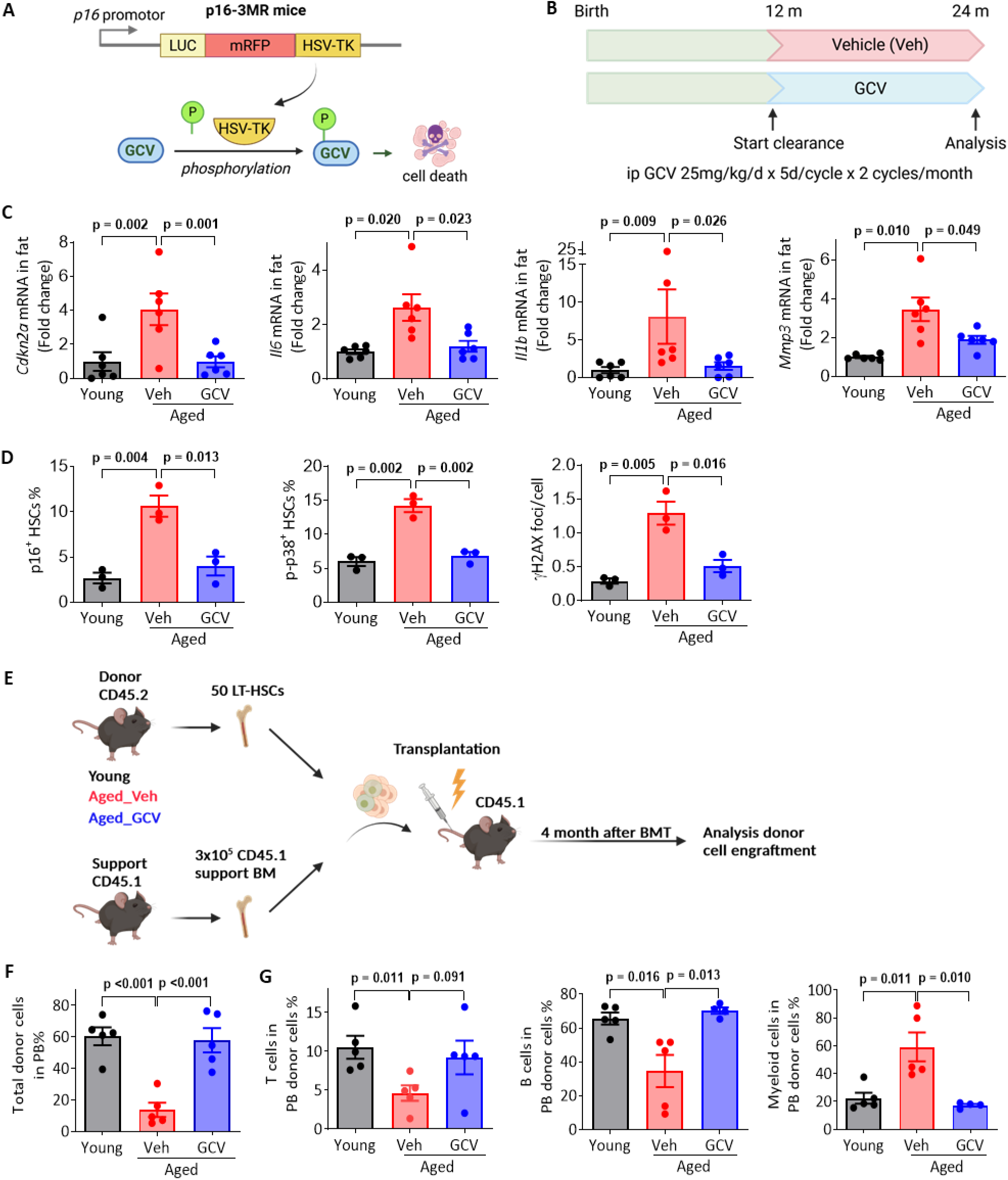
Clearance of senescent cells in aged p16-3MR mice reverses myeloid bias in hematopoiesis. (A) Schematic of the p16-3MR transgene and strategy used to selectively clear senescent cells. (B) p16-3MR mice were treated starting at 12 months of age with PBS (Aged-Veh) or GCV (Aged-GCV) for an additional 12 months before analysis. (C) Expressions levels of key senescence and inflammatory marker genes in adipose tissue, assessed by qRT-PCR; mean ± SEM, Student’s t-test. (D) Percentage of cells with detectable p16, phosphorylated p38, or multiple γH2A.X foci measured by immunofluorescence staining. (E) Experimental outline for competitive BMT. (F-G) Donor chimerism (F) and lineage distribution (G) in peripheral blood was assessed by flow cytometry; mean ± SEM, Student’s *t*-test.

**SFigure 1.**
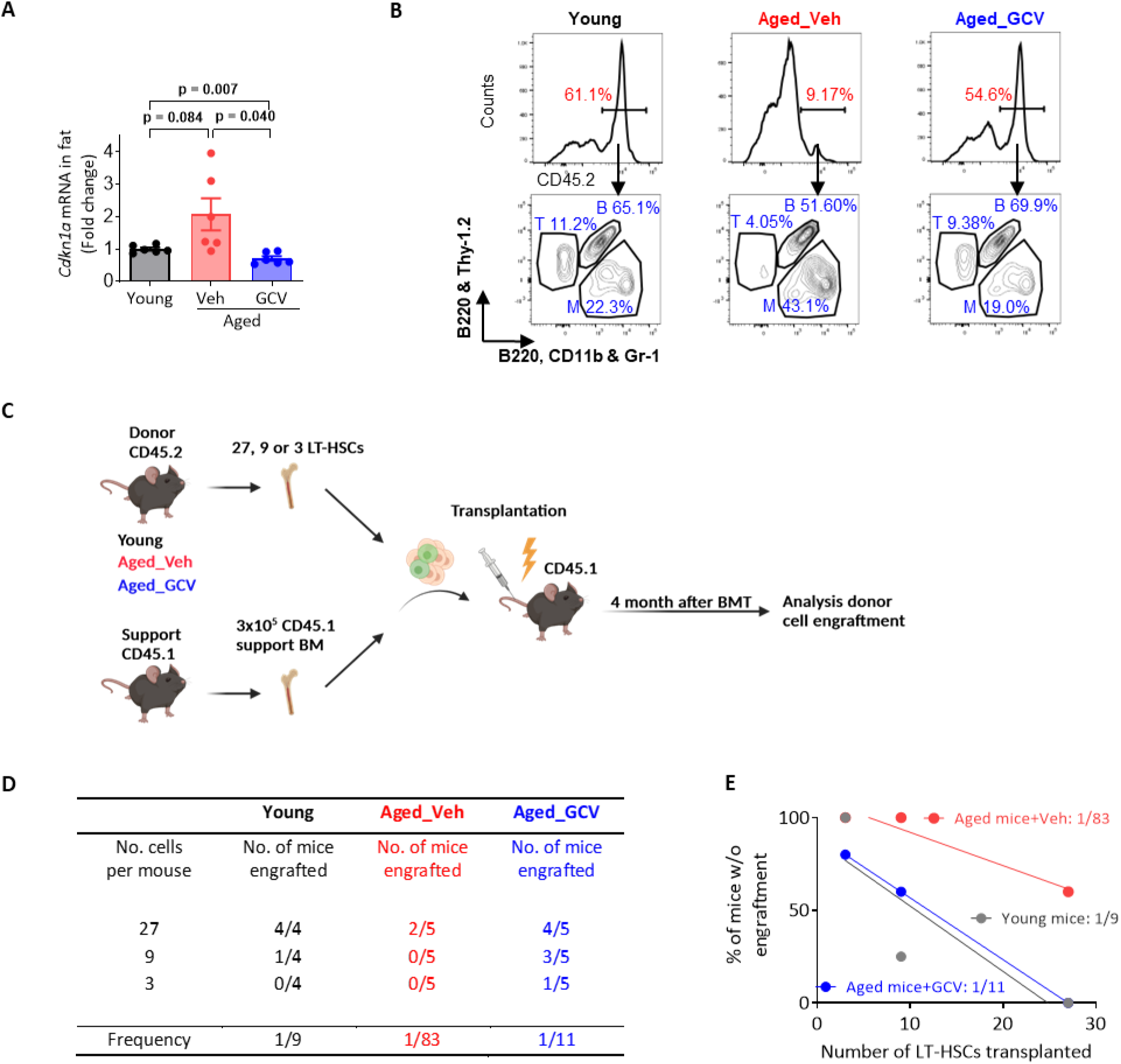
Clearance of senescent cells mitigates HSC functional decline in naturally aged p16-3MR mice. (A) *Cdkn1a* transcript levels in adipose tissue (qRT-PCR); mean ± SEM, Student’s *t*-test with Welch’s correction. (B) Flow cytometry testing donor chimerism (CD45.2%) and peripheral blood lineage composition (T cells, B cells, myeloid cells) at 4 months after BMT using donor material from young, aged vehicle-treated, or aged GCV-treated mice. (C-E) Design of the limiting dilution transplantation assay (C), frequency of reconstitution (D), and negative binomial regression analysis of the percentage of non-engrafted mice versus number of immunophenotypic LT-HSCs transplanted (E).

Collectively, these results demonstrate that clearance of p16-expressing senescent cells in aged mice reduces senescence-associated inflammation and DNA damage, correcting age-associated myeloid lineage bias and preserving regenerative function of the HSC population.

### Clearance of senescent cells rejuvenates BM mesenchymal stromal cells and restores hematopoietic support

To assess the impact of senescent cell clearance on the bone marrow microenvironment, we analyzed gene expression and functional properties of mesenchymal stromal cells (MSCs) isolated from young, aged vehicle-treated, and aged GCV-treated p16-3MR mice. Quantitative RT-PCR revealed elevated expression of pro-inflammatory and senescence-associated genes, including *Ccl5*, *Il1β*, and *Tnfα*, in aged vehicle-treated MSCs compared to young controls (Fig. 2A). In contrast, GCV administration led to significantly reduced expression of these SASP genes compared to aged vehicle-treated animals, indicating effective senescent cell clearance and suppression of MSC-mediated inflammation. Next, we assessed the functional ability of MSCs to support hematopoietic stem and progenitor cell (HSPC) growth. FACS-sorted Lin^-^Sca-1^+^c-Kit^+^ (LSK) HSPCs from young, aged vehicle-treated, and aged GCV-treated mice were co-cultured with MSCs (CD45^-^Lin^-^PI^-^) from each of the 3 groups grown as a monolayer over 3 days (Fig. 2B) followed by colony-forming unit (CFU) assay. Young LSK HSPCs cultured atop aged vehicle-treated MSCs yielded significantly fewer multipotent CFU-GEMM (granulocyte/erythroid/monocyte/megakaryocyte) and lineage-restricted CFU-GM (granulocyte/macrophage) colonies compared to cultures with young MSCs, reflecting the functional decline of the aged bone marrow niche (Fig. 2C). Notably, aged GCV-treated MSCs supported substantially higher numbers of both myeloid and multilineage colonies, comparable to the young MSC. Similar restoration of colony-forming potential was observed when aged GCV-treated LSK cells were co-cultured with aged GCV-treated MSCs versus vehicle controls. These findings show that clearance of senescent cells in aged p16-3MR mice reverses the inflammatory phenotype of MSCs and enhances their capacity to support HSPC function *ex vivo*, thereby contributing to rejuvenation of the BM niche and by extension of hematopoiesis as a whole.

**Figure 2.**
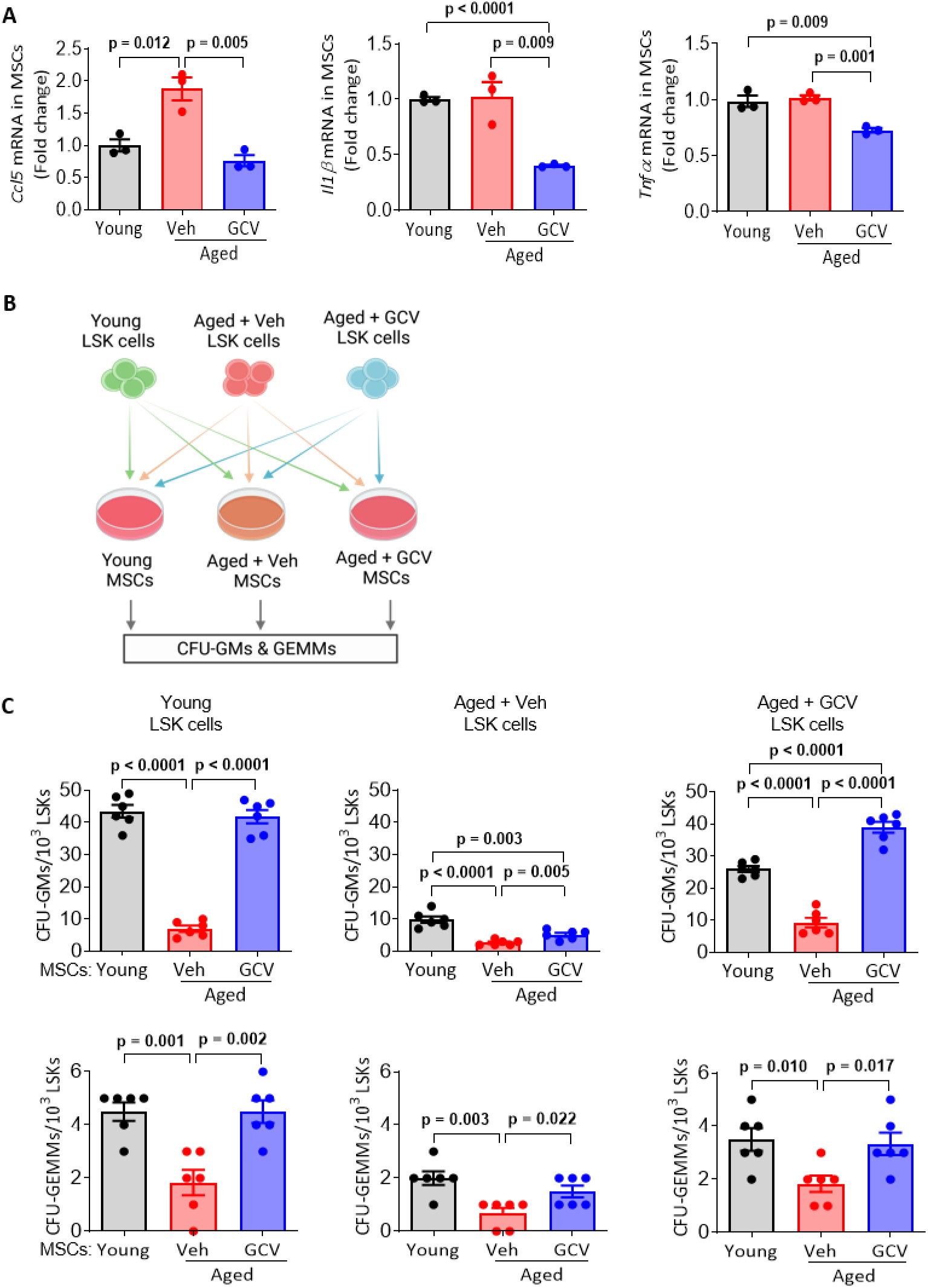
Clearance of senescent cells improves the functional ability of BM MSC to support hematopoiesis. (A) Quantitative RT-PCR analysis of gene expression levels in BM MSCs from young, aged vehicle-treated, and aged GCV-treated p16-3MR mice; mean ± SEM, Student’s *t*-test. (B) Schematic of the *ex vivo* CFU assay in co-culture with MSCs isolated from each of the 3 experimental groups. (C) CFU-GM and CFU-GEMM output in indicated LSK/MSC combinations; mean ± SEM, Student’s *t*-test.

### 753b exhibits efficient, selective senolytic activity with low hematologic toxicity in aged animals

Our findings in p16-3MR mice provide a compelling proof-of-concept that eliminating p16⁺ senescent cells ameliorates age-related hematopoietic decline. Yet, this model is translationally limited by a non-physiologic transgene and requirement for GCV administration. Studies relying on existing senolytic drugs such as dasatinib plus quercetin or the BCL-2 family inhibitors ABT-737 and navitoclax demonstrated that pharmacologic senolysis is feasible in principle^20,23–25^. Despite their promise, these approaches are complicated by suboptimal selectivity, bioavailability and dose-limiting hematologic toxicities, including thrombocytopenia and neutropenia, which hinder long-term treatment necessary for efficacy^32^.

753b is a BCL-2/BCL-XL PROTAC that recruits both BCL-XL and BCL-2 to the VHL E3 ligase, promoting their ubiquitination and degradation selectively in VHL-expressing cells. As the result, 753b spares platelets, which express very low levels of VHL and thus is less toxic to platelets than navitoclax (ABT263)^29,31^. These features provide a strong rationale to evaluate 753b as a next-generation senolytic agent with an improved hematologic safety profile in aged animals.

As shown in our recent reported studies^30^, 753b exhibited high selective cytotoxicity against irradiation-induced senescent cells (IR-SCs) in WI-38 human fibroblasts, with an EC_50_ of approximately 0.134 µM, whereas confluent and sub-confluent non-senescent control WI-38 cells showed minimal loss of viability over the same concentration range (SFig. 2A). Pharmacokinetic analysis in mice demonstrated that a single 5 mg/kg intraperitoneal (IP) injection of 753b yielded measurable plasma levels with sustained exposure over time, suggesting a favorable *in vivo* stability profile (SFig. 2B). Next, we evaluated hematologic safety of 753b in aged mice. Body weights of vehicle- and 753b-treated animals were comparable in both males and females, suggesting no overt systemic toxicity under this dosing regimen (SFig. 2C). Further, red blood cell counts were indistinguishable between aged vehicle- and 753b-treated mice in both sexes following 8 doses of treatment, indicating no major adverse effect on erythropoiesis (SFig. 2D). Aged vehicle-treated mice exhibited elevated platelet counts compared to young controls consistent with age-associated thrombocytosis, whereas 753b-treated aged animals displayed reduced platelet counts similar to levels characteristic of young mice without thrombocytopenia (SFig. 2E), implying correction of an age-related hematologic abnormality rather than platelet toxicity or myelosuppression. Finally, we found increased expression levels of the senescence marker gene *Cdkn2a* (*p16*) in the spleens of aged vehicle-treated mice compared to young control animals, consistent with an elevated senescent cell burden. At the same time, 753b treatment significantly reduced *Cdkn2a* expression in aged animals, in alignment with senescent cell clearance (SFig. 2F). Together, these data demonstrate that 753b is an effective and selective senolytic *in vitro* and *in vivo*, significantly reducing expression of senescence marker genes while maintaining favorable pharmacokinetics and a low toxicity profile in aged animals.

**SFigure 2.**
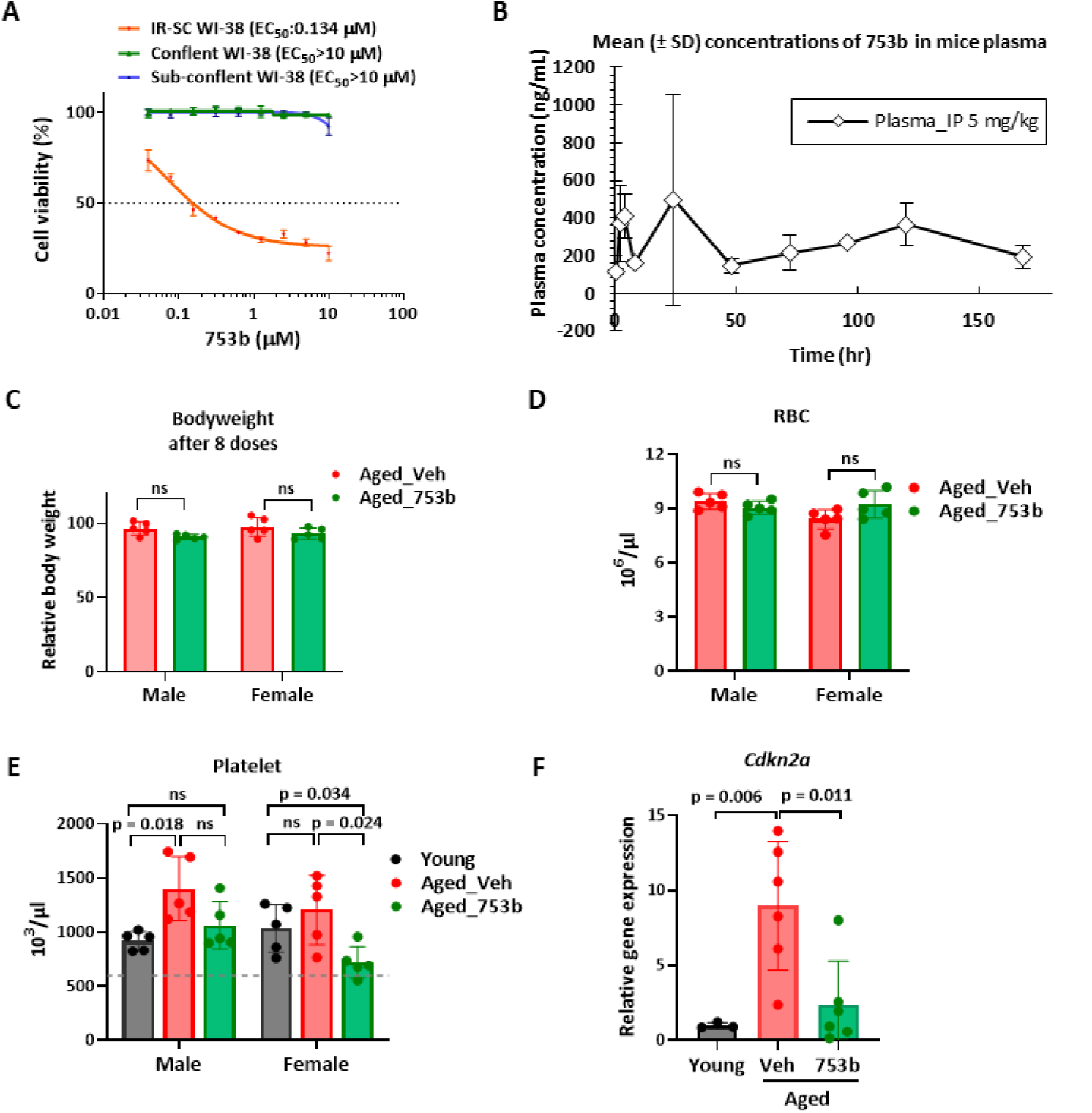
753b targets senescent cells with high efficacy and low toxicity. (A) Dose-response curves to 753b in confluent, sub-confluent and senescent (IR-SC) WI-38 cells (IR-SC) after 72 hours of treatment; mean ± SD. (B) Plasma concentration of 753b in mice following a single IP injection (5 mg/kg). Plasma samples were measured at 0.5, 2, 4, 8, 24, 48, 72, 96, 120 and 168 hours post injection; mean ± SD. (C) Body weight of aged mice after 8 doses of 753b or vehicle in both sexes; mean ± SEM; Student’s *t*-test. (D) Red blood cell (RBC) counts in peripheral blood of aged male and female mice treated with vehicle or 753b; mean ± SEM; Student’s *t*-test. (E) Platelet counts in young, aged vehicle-treated, and aged 753b-treated mice; mean ± SEM; Student’s *t*-test. (F) Relative gene expression of *Cdkn2a* (coding for p16) in spleens of young, aged vehicle-treated, and aged 753b-treated mice; mean ± SEM; Student’s *t*-test with Welch’s correction.

### 753b-mediated senescent cell clearance reverses myeloid bias in aged HSCs

Myeloid skewing is a hallmark of aged HSCs, characterized by predominant myeloid output and reduced lymphoid production that together drive immune decline and increased susceptibility to myeloid malignancies. To determine whether 753b-mediated senescent cell clearance functionally restored more balanced mature cell output of HSCs, BMT assays were performed using equal numbers of donor cells (CD45.2⁺) from 100-week-old mice treated with vehicle (Aged_Veh) or 753b (Aged_753b) for 3 weeks and from young controls that were mixed with congenic CD45.1^+^ support and grafted into lethally-irradiated recipients (Fig. 3A). Compared to recipients of young BM controls, mice implanted with vehicle-treated aged BM displayed a markedly increased proportion of myeloid cells in peripheral blood at the expense of B-cell output in the donor CD45.2⁺ graft, at both 4 and 12 weeks after transplantation as expected (Fig. 3B-C, SFig. 3A-B). In contrast, pre-treatment with 753b led to a significant reversal of the myeloid bias in aged BM evidenced by a more balanced lineage output similar to that of young donors. The CD45.1⁺ support compartment as an internal control showed no significant differences in lineage distribution among groups, confirming that the observed changes were intrinsic to the donor HSC compartment (SFig. 3C). Analysis of HSPC populations further aligned with a functional rejuvenation of aged HSCs after 753b treatment. HSPCs from Aged_Veh donors showed increased proportion of granulocyte-macrophage progenitors (GMP) with a relative decrease of megakaryocyte-erythroid progenitors (MEP), consistent with myeloid-biased differentiation. These changes were reversed in Aged_753b group, resembling progenitor composition seen in mice engrafted with BM from young donors (Fig. 3D-E). Similarly, myeloid progenitor distribution within the internal control CD45.1⁺ compartment remained largely unchanged across groups, reinforcing the notion of self-intrinsic alterations to lineage output in the donor BM (SFig. 3D). Consistently, CFU assay found a trend toward increased CFU-GM in the Aged_Veh group and partial rescue after 753b treatment (SFig. 3E). Collectively, these findings demonstrate that pharmacologic clearance of senescent cells with 753b alleviates myeloid expansion and restores a more balanced hematopoietic output from aged HSCs, suggestive of rejuvenation.

**Figure 3.**
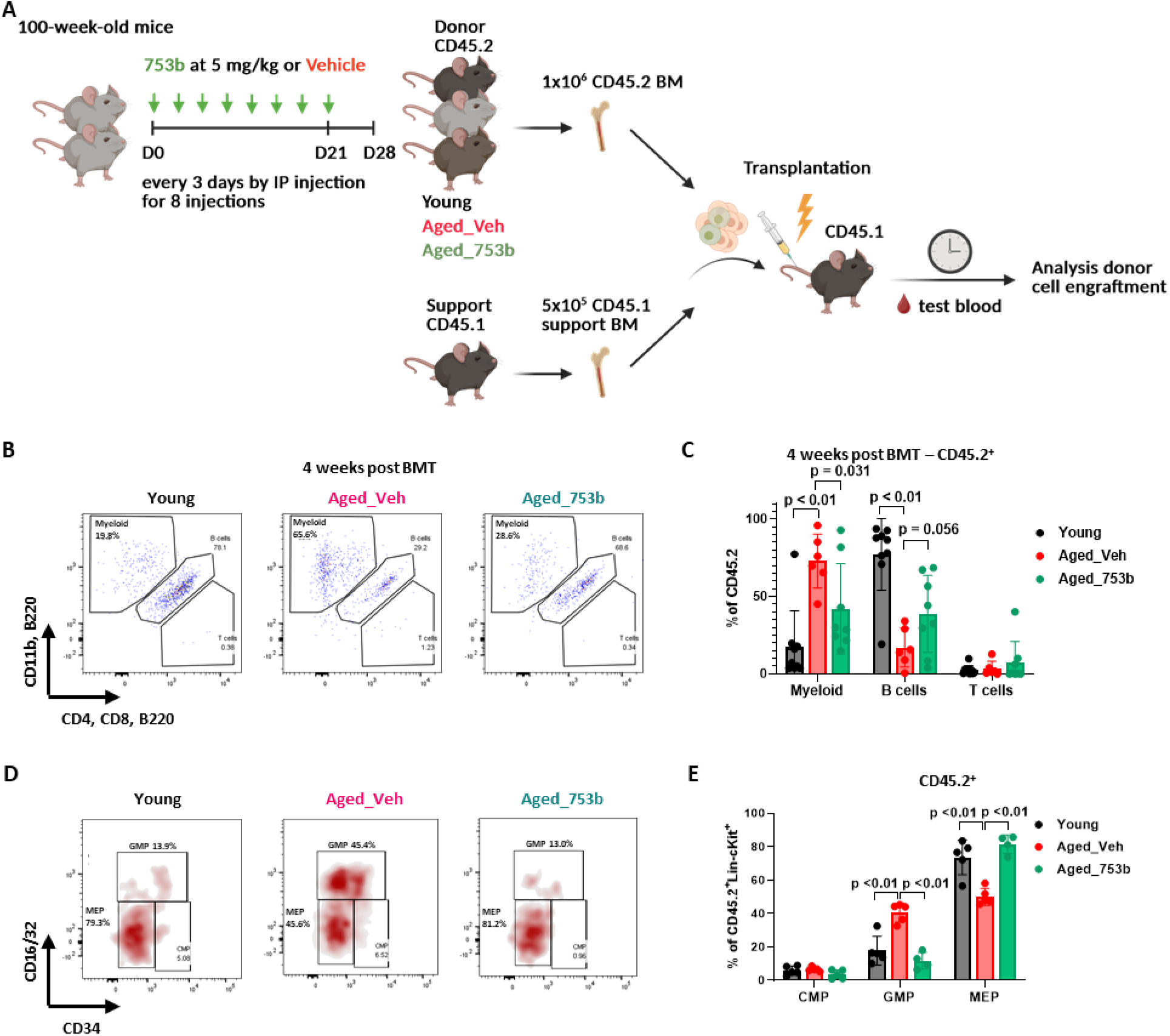
Clearance of senescent cells with 753b reverses myeloid bias in aged HSCs. (A) Schematic of the experimental workflow. (B, C) Representative flow cytometry plots of peripheral blood leukocytes 4 weeks after BMT (B), with quantification of CD45.2^+^ donor-derived myeloid, B, and T cell output (C); mean ± SEM; Student’s *t*-test. (D) Flow cytometry analysis of bone marrow progenitor composition in CD45.2^+^ LSK population. (E) Frequencies of CMP (common myeloid progenitor), GMP (granulocyte-macrophage progenitor) and MEP (megakaryocyte-erythroid progenitor) populations in indicated groups; mean ± SEM; Student’s *t*-test.

**SFigure 3.**
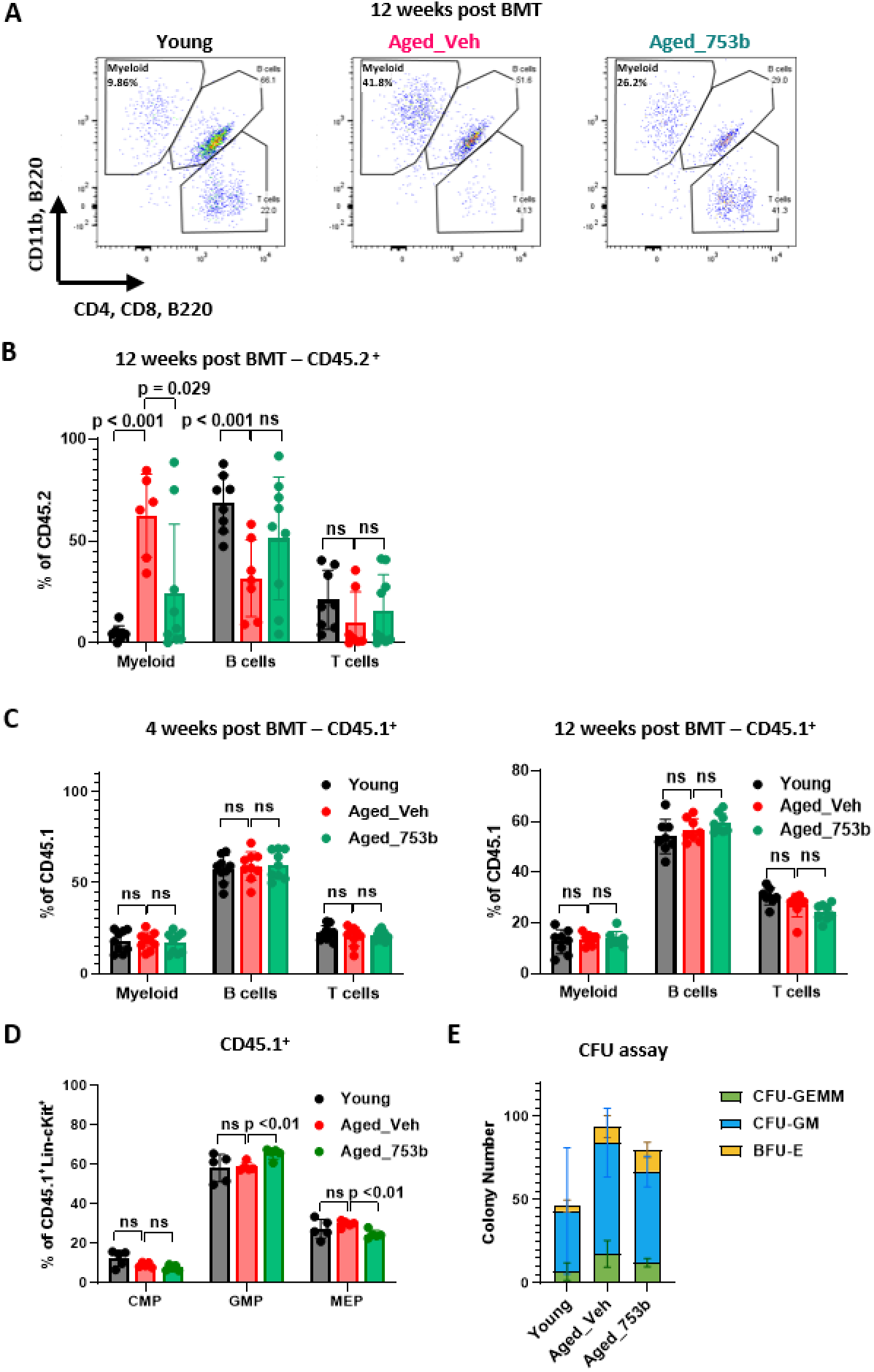
Clearance of senescent cells with 753b reverses myeloid expansion contributed by aged HSCs. (A, B) Representative flow cytometry plots of peripheral blood leukocytes 12 weeks after BMT (A) and quantification of myeloid, B, and T cells in CD45.2^+^ donor-derived compartment (B). (C) Myeloid, B, and T cell percentages in wild-type control competitor-derived CD45.1^+^ compartment 4 and 12 weeks post-BMT; mean ± SEM; Student’s *t*-test. (D) Frequencies of CMP, GMP and MEP populations in competitor-derived CD45.1^+^ compartment; mean ± SEM; Student’s *t*-test. (E) CFU assay showing the numbers of colony types across indicated groups; mean ± SD.

### Clearance of senescent cells with 753b reshapes transcriptionally-defined composition of aged HSCs toward a more youthful state

To gain mechanistic insight into 753b effects on aged HSCs, we profiled single-cell transcriptomes from a total of 34,000 FACS-purified HSCs (LSK CD150⁺) from young, aged vehicle-treated (Aged_Veh), and aged 753b-treated (Aged_753b) mice using the 10x Genomics Chromium platform (Fig. 4A). Unsupervised clustering using Seurat package identified 19 transcriptionally distinct clusters that were annotated based on established marker genes^33^ (Fig. 4B, SFig. 4A) and further merged along the differentiation hierarchy into non-primed (stem, 6 clusters), intermediate (5 clusters), and primed (lineage-progenitor, 8 clusters) subgroups (Fig. 4B). Consistent with previous observations, aged HSCs showed a pronounced expansion of non-primed populations with a reduction of primed clusters (Fig. 4B-C). Interestingly, in Aged_Veh animals, elevated *Bcl2* expression was primarily observed in np1 and ifn clusters (long-term and short-term HSCs (LT-HSCs and ST-HSCs)) whereas the tgf cluster (mainly LT-HSCs) preferentially upregulated *Bcl2l1* (encoding Bcl-xL) (SFig. 4B), suggesting engagement of distinct pro-survival signaling mechanisms among HSC subpopulations and underscoring the importance of dual targeting of BCL-2 and BCL-xL. Notably, a short 8-dose course of 753b treatment markedly reduced *Bcl2/Bcl2l1* transcript levels in HSCs, consistent with depletion of senescent HSCs (SFig. 4B). 753b treatment considerably rebalanced aging-related changes in stem and progenitor cell distribution in that the proportion of non-primed HSCs decreased, while the relative abundance of cells in intermediate clusters increased along with slight elevation of primed progenitors (Fig. 4B-C, SFig. 4C). Trajectory and pseudotime analyses further supported partial rejuvenation of aged HSCs by 753b (Fig. 4D). Aged_Veh HSCs were enriched earlier in pseudotime, whereas Aged_753b HSCs were more distributed along pseudotime differentiation paths, indicating restoration of a more progressive, stem-to-progenitor differentiation continuum resembling that of young HSCs. Cell-cycle analysis showed that baseline proliferation was low in non-primed clusters, with Aged_Veh HSCs exhibiting further reductions in cycling cells (Fig. 4E, SFig. 4D). In contrast, 753b treatment produced consistent, albeit modest, increases in cycling (fraction in S/G2/M phases) within all non-primed stem populations including tgf, np1, and np2 clusters (Fig. 4E). These changes indicate partial restoration of proliferative competence within the stem cell compartment of aged mice after 753b treatment. Intermediate and primed clusters displayed similar cycling patterns across all groups, suggesting that aging-related defects in proliferation and differentiation predominantly affect the most primitive HSCs (SFig. 4D).

**Figure 4.**
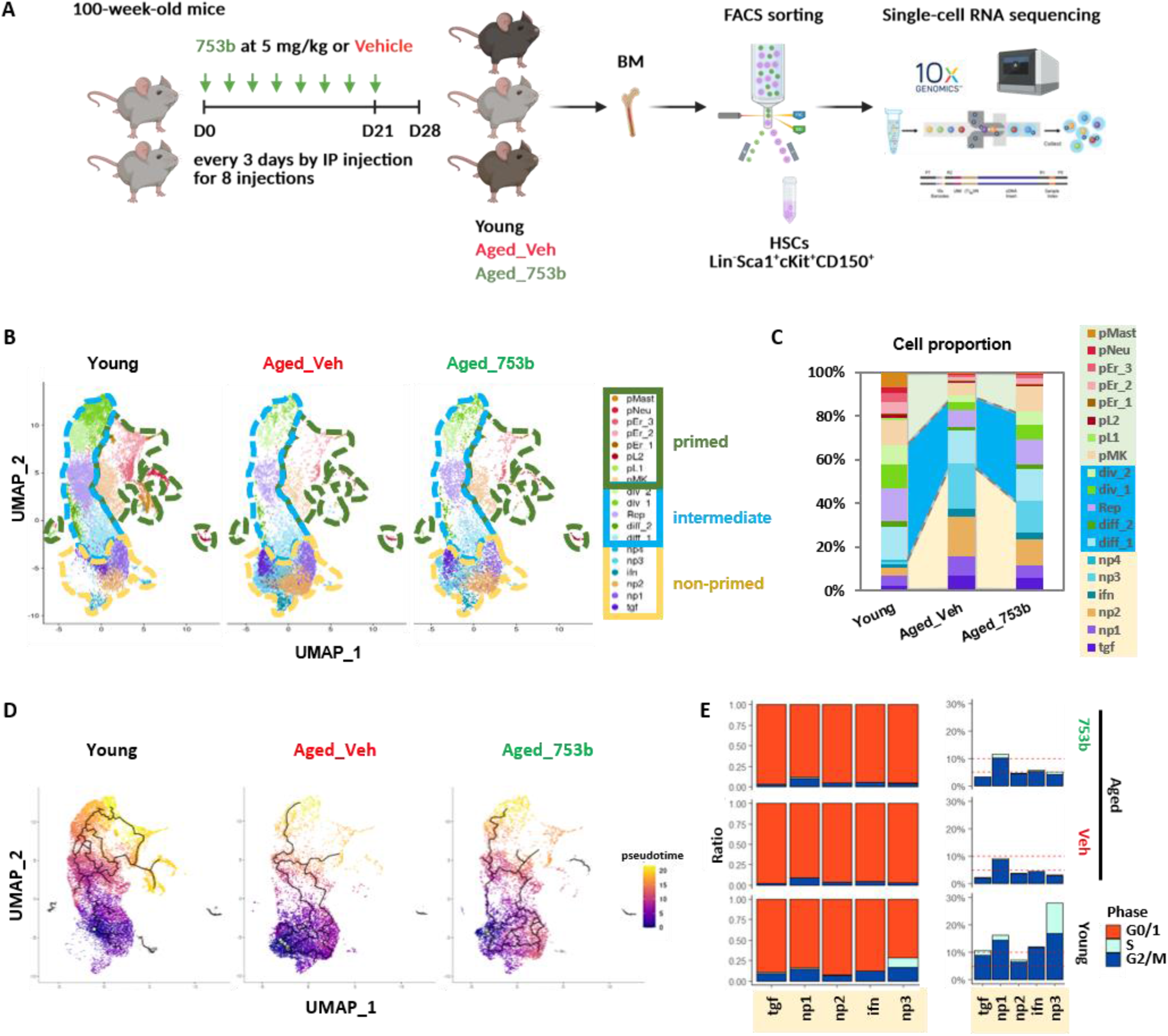
Clearance of senescent cells with 753b rejuvenates the transcriptomes of aged HSCs. (A) Sample preparation for scRNA-sequencing using BM from young, aged vehicle-treated, and aged 753b-treated mice. Pools of FACS-sorted LSK CD150^+^ HSCs from both male and female animals (n≥6 per group) were processed using 10X Genomics platform. (B) A UMAP projection showing the distribution of the 19 transcriptionally-defined HSC clusters, subgrouped into non-primed (yellow), intermediate (blue) and primed (green) metaclusters. (C) Proportion of cells in each cluster by age and treatment type. (D) HSC differentiation trajectory in pseudo-time. (E) Ratio of cell cycle (G1, S and G2/M) phases in non-primed clusters in indicated groups.

**SFigure 4.**
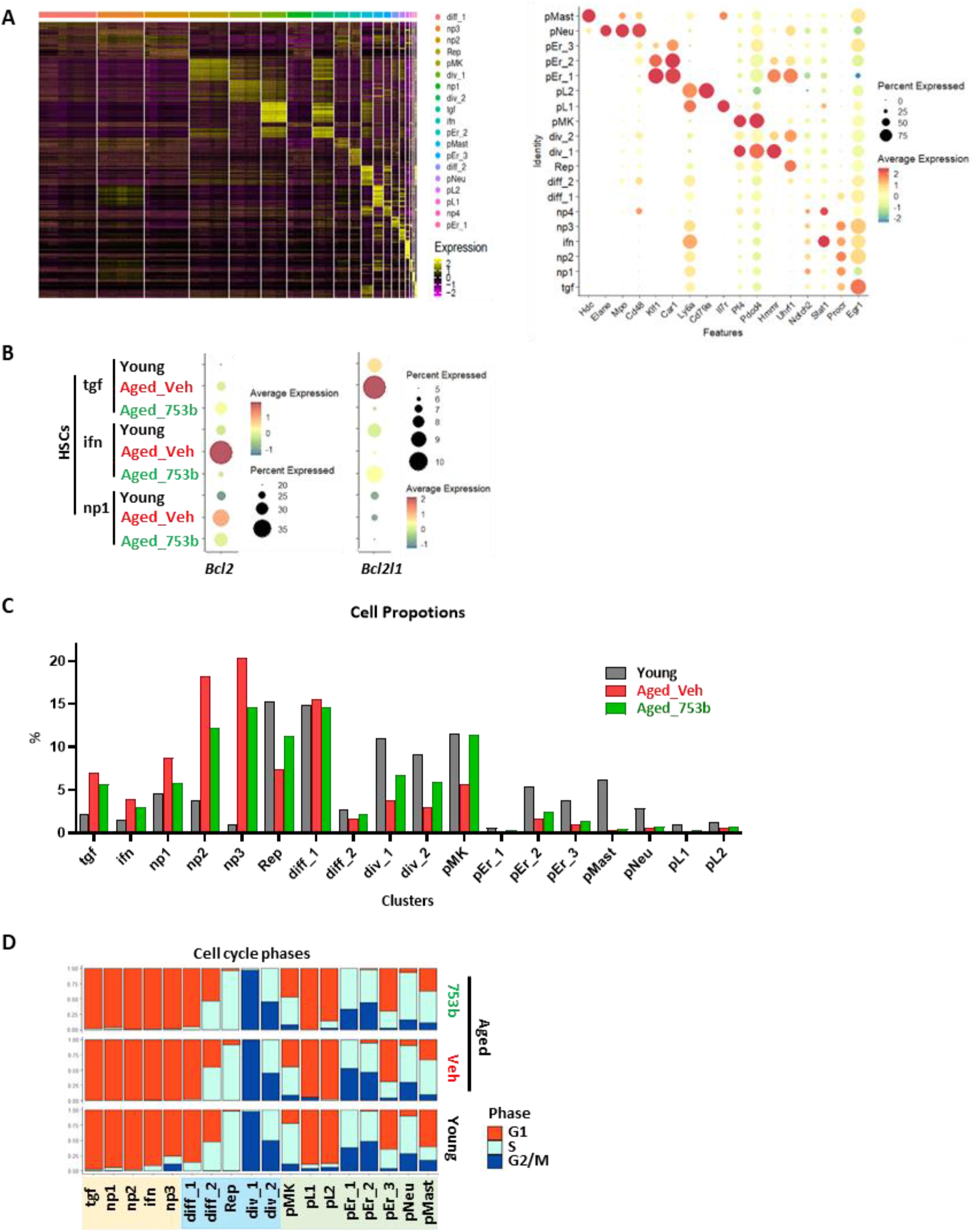
Clearance of senescent cells with 753b reshapes the transcriptomes of aged HSCs towards a more youthful composition of subpopulations. (A) Left, heatmap of scaled expression values for the top marker genes defining each cell cluster identified in single-cell RNA seq data. Right, dot plot showing representative cluster-specific marker gene expression levels, where dot size represents the percentage of cells expressing each gene and color indicates average expression level. (B) *Bcl2* and *Bcl2l1* (*Bcl-xL*) expression levels in tgf, ifn, and np1 HSC clusters across conditions. (C) Cluster proportions across conditions. (D) Distribution of cell cycle phases (G1, S, G2/M) across clusters and conditions.

Together, these analyses indicate that 753b-mediated senescent cell clearance rejuvenates the aged HSC compartment, normalizes the balance between stem-like and progenitor populations, and partially restores proliferation and repopulation capacity of aged HSCs.

### 753b treatment attenuates aging-associated transcriptional signatures in HSCs

To quantify aging-associated transcriptional alterations in HSCs, we defined an aging signature using genes with most significantly increased expression in aged non-primed HSC clusters (tgf, np1, np2, ifn, np3) that encompass LT-HSCs and ST-HSCs. Importantly, a subset of genes in our signature, *Cavin2*, *Clu*, *Gadd45g*, *Mt1*, *Nrgn*, *Nupr1*, *Plscr2*, *Prtn3*, *Rorb*, *Selp*, and *Sult1a1*, were previously reported as highly expressed in aged HSCs^2,34,35^ (Fig. 5A, red). Among those, we found significant enrichment genes involved in inflammatory signaling, oxidative stress response, metabolic, adhesion, and apoptotic pathways characteristic of aged HSCs (Fig. 5B). While a large subset of these genes showed robust upregulation in aged HSCs, their expression was markedly reduced upon 753b treatment, indicating partial reversal of the aging-associated transcriptional state (Fig. 5C). Next, we evaluated the utility of our gene signature (Fig. 5A) for capturing conserved aging features in previously published independent bulk RNA-seq datasets from murine and human HSCs^1,36^, using AddModuleScore function in Seurat^37^ to compute aging scores (SFig. 5). The composite aging score was significantly higher in sorted LSK CD48⁻CD150⁺ HSCs from old mice (20-24 months) than in young (2-3 months) controls (SFig. 5A), indicating robust discrimination of physiologic HSC aging in an independent validation cohort (SFig. 5A). Similarly, aging scores were significantly elevated in human Lin⁻CD34⁺CD38⁻ HSCs from aged (65-75 years) donors compared to young (18-30 years) individuals (SFig. 5B), demonstrating cross–species conservation of the aging–associated transcriptional programs. Albeit aging–score genes showed marked expression heterogeneity in human samples, these analyses validate our aging gene signature as a quantitative metric of aging across murine and human HSC transcriptomes.

**Figure 5.**
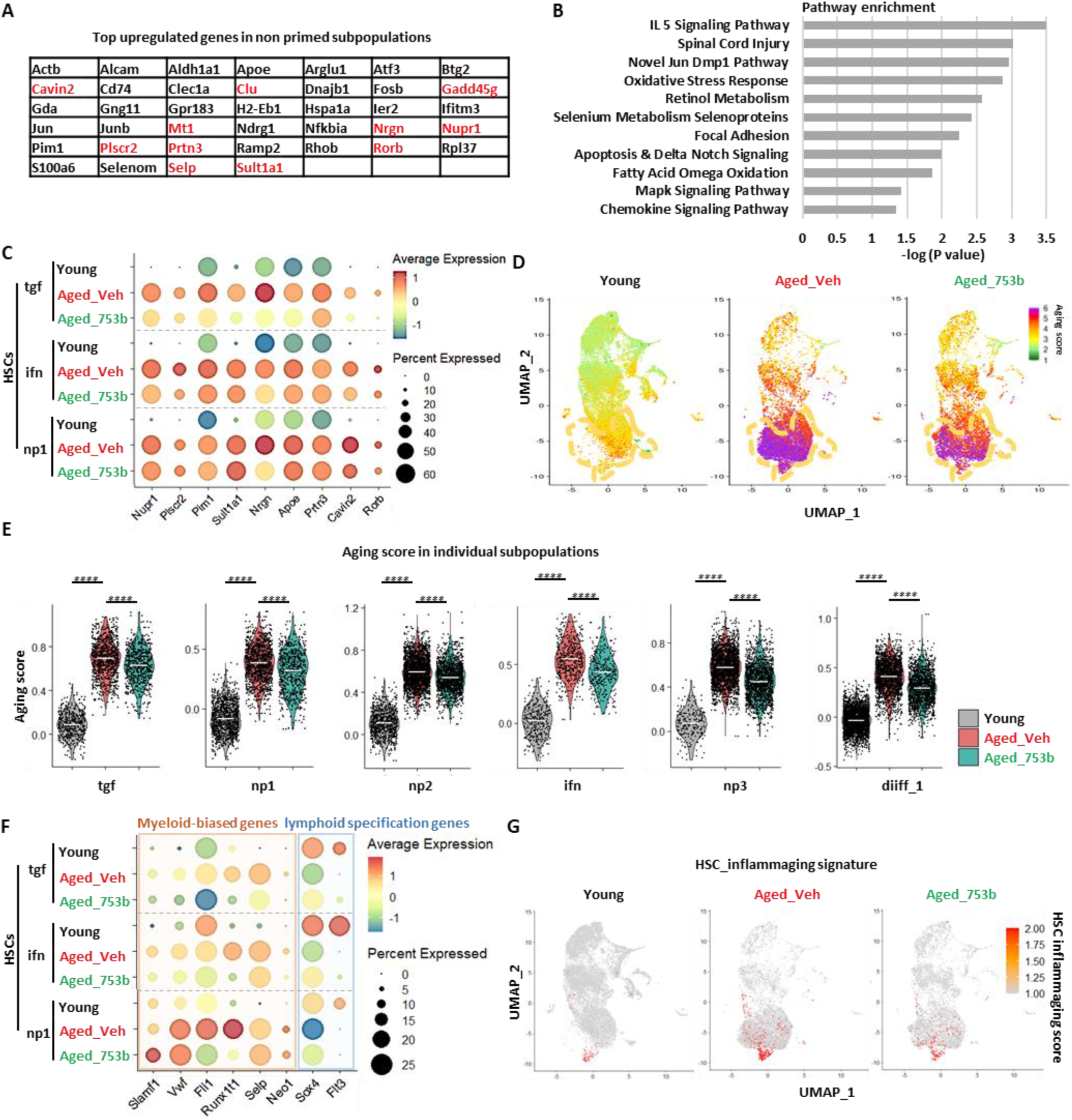
753b treatment ameliorates aging–associated transcriptional features in hematopoietic stem cells. (A-C) Top genes elevated in aged non-primed HSCs (A), most significantly enriched pathways these genes contribute to (B), and their expression metrics in indicated groups (C). (D) Aging scores calculated based on expression levels of genes listed in (A). A UMAP projection of all cells color-coded according to aging score for each condition. Yellow dashed outline indicates non–primed HSCs. (E) Aging scores for cells within individual clusters, horizontal lines denote the median, Wilcoxon rank-sum test. (F) Expression of myeloid-biased and lymphoid specification genes in HSCs from young, aged vehicle-treated (Veh), and aged 753b-treated mice. (G) A UMAP projection of all cells color-coded to reflect inflammaging score for each condition.

**SFigure 5.**
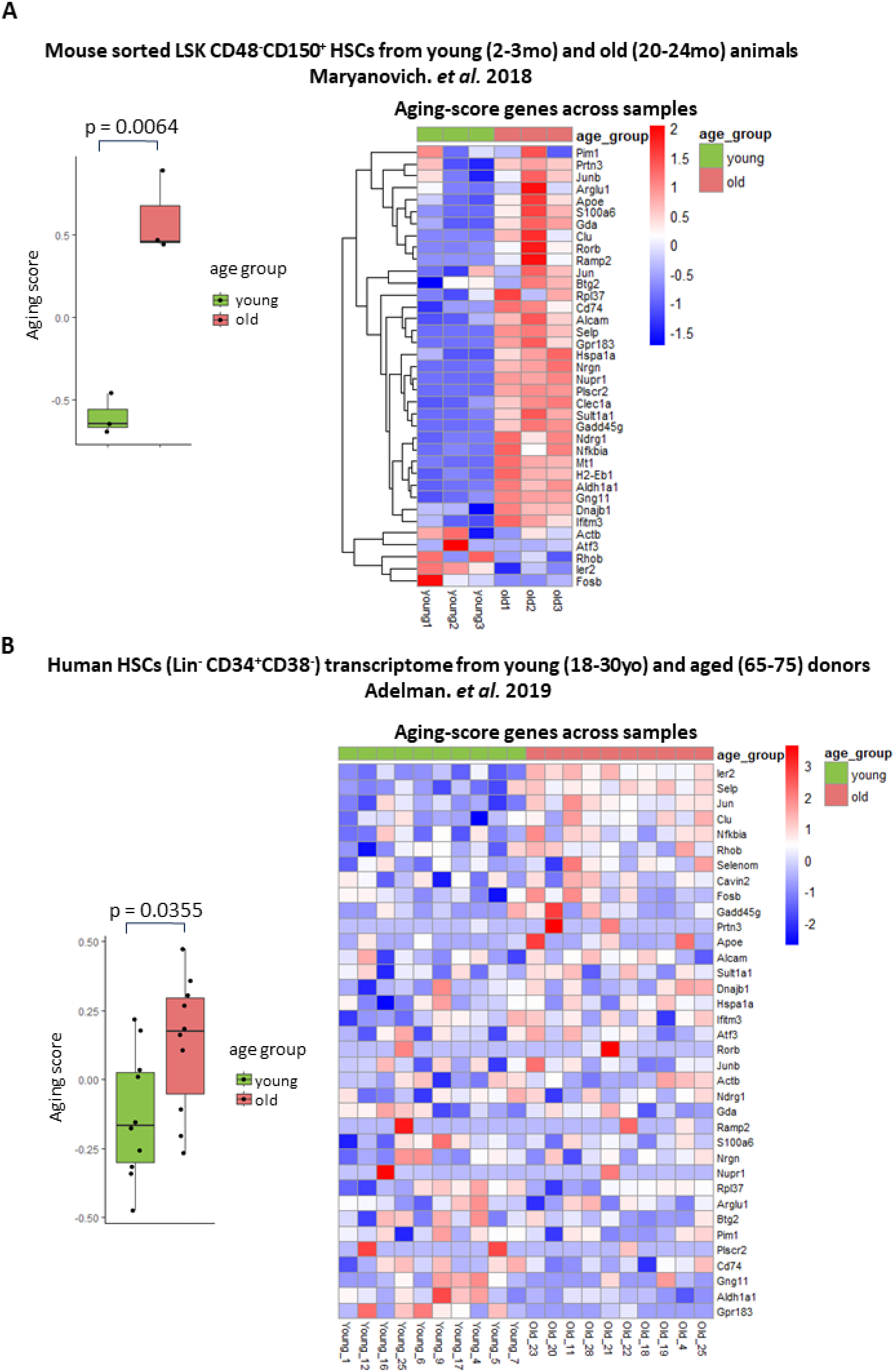
Consolidating aging score in independent mouse and human HSCs datasets. Boxplot (left) comparing age groups using the aging score derived from our single-cell–based aging gene set, calculated for published bulk transcriptomes of sorted murine LSK CD48^-^CD150^+^ HSCs from young (2-3mo) and old (20-24mo) mice (A) and human HSC (Lin^-^CD34^+^CD38^-^) transcriptomes from young (18-30yo) and aged (65-75) donors (B). Heatmap (right) shows z-scaled expression of aging-score genes across samples with annotation bars indicating age groups.

**SFigure 6.**
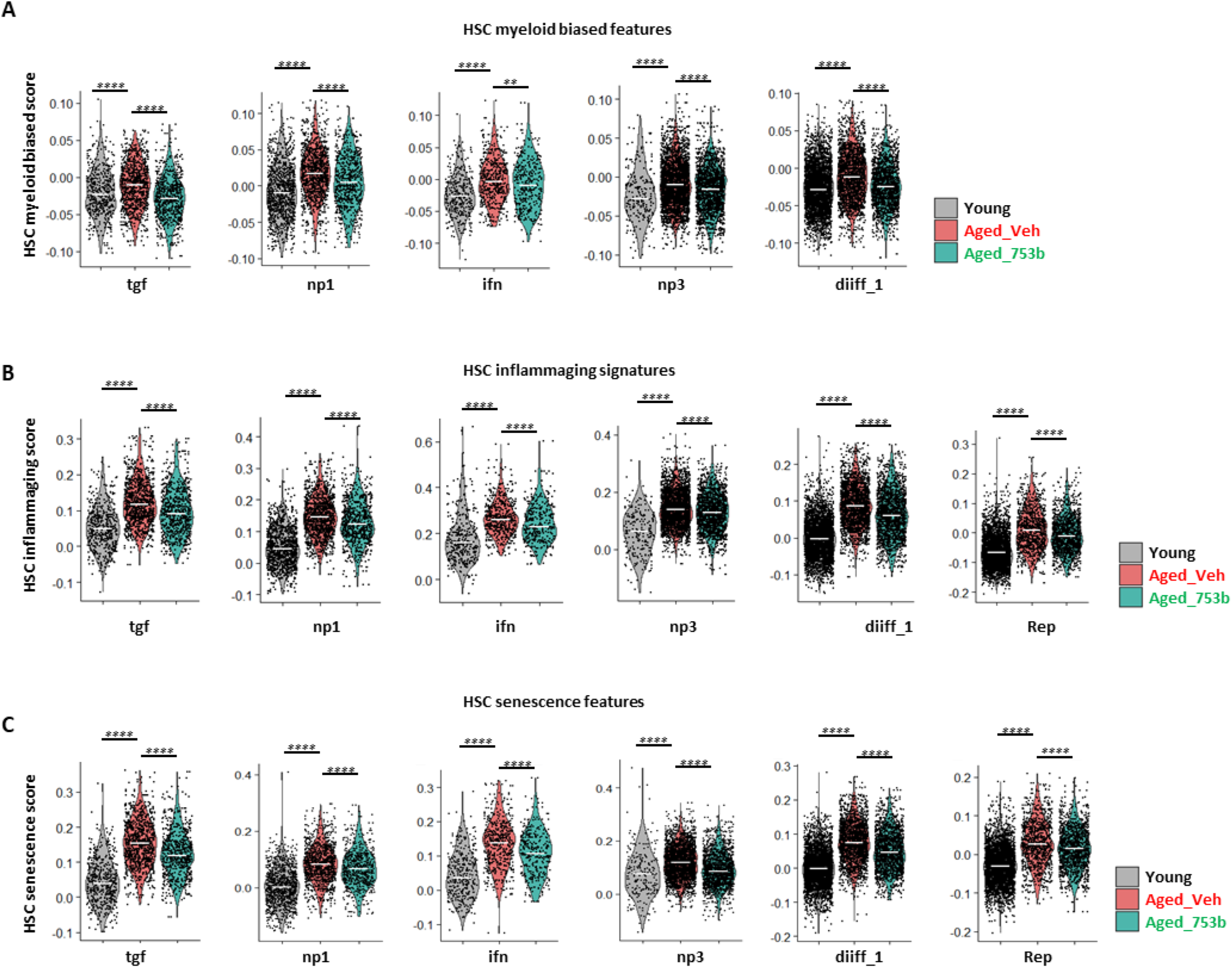
753b treatment ameliorates aging–associated transcriptomic features in HSCs. (A–C) Gene signature scores for HSC myeloid-biased features (A), HSC inflammaging signatures (B), and HSC senescence features (C) across the indicated clusters in young, aged_veh and aged_753b mice. Each point represents a single cell; violin width reflects the distribution of scores within each group, and horizontal lines denote the median, Wilcoxon rank-sum test.

To gain a granular view of aging-related changes to gene expression within the HSPC compartment, we calculated aging scores of individual single-cell transcriptomes and projected onto UMAPs (Fig. 5D). Aged vehicle-treated HSCs exhibited high aging scores, particularly within the most primitive non-primed compartment, whereas 753b treatment markedly reduced the density of high-aging-score cells, suggesting partial transcriptional rejuvenation of HSCs (Fig. 5D). Analysis restricted to non-primed clusters found that aging scores were consistently elevated in all aged HSCs but showed significant reductions following 753b exposure, indicating that senescent cell clearance rejuvenates multiple HSC compartments (Fig. 5E). Collectively, these transcriptional changes, quantitatively measured using our scoring approach, support a notion that 753b-mediated senescent cell clearance rejuvenates aged HSCs at the molecular level, attenuates aging-score gene expression across multiple non-primed HSC clusters, ameliorating key features of HSC aging.

### Senescent cell clearance by 753b reverses myeloid bias within aged HSCs, attenuates inflammaging and senescence signatures

Age–associated HSC dysfunction is characterized by skewed lineage output and chronic inflammatory signaling that together drive ineffective hematopoiesis. To determine whether 753b–mediated senescent cell clearance can correct these functional hallmarks of HSC aging, we next examined gene expression features related to myeloid bias, inflammaging, and senescence in aged HSCs. Aged vehicle-treated HSCs were characterized by elevated expression of myeloid lineage genes (*Slamf1, Vwf, Fli1, Runx1t1 Sell, Neo1*) and reduced expression of lymphoid specification factors (*Sox4, Flt3*) compared with young HSCs (Fig. 5F). Gene-signature scores for HSC myeloid-biased features^38,39^ were also significantly higher in non-primed clusters in aged vehicle-treated HSCs (SFig. 6A). Senolytic 753b treatment markedly reduced myeloid-biased signatures while partially restoring expression of lymphoid-associated genes (Fig. 5F, SFig. 6A). Similarly, aged_Veh HSCs exhibited elevated inflammaging and senescence signatures^39^ relative to young HSCs (Fig. 5G, SFig. 6B-C), reflecting chronic inflammatory signaling and activation of stress–response programs characteristic of aged stem cells. In contrast, these transcriptional signatures were markedly attenuated by 753b treatment, with distributions shifting toward those of young HSCs in most clusters (Fig. 5G; SFig. 6B-C). Taken together, these functional and single-cell gene expression studies show that BCL-xL and BCL-2 targeted pharmacological clearance of senescent cells with 753b reverses myeloid bias, enhances lymphoid potential, and restores more balanced hematopoietic differentiation of aged HSCs, providing a mechanistic basis for the improved repopulation capacity after transplantation *in vivo*.

### Short-term 753b treatment achieves targeted senolysis with preserved bone marrow stromal cell composition and cycling

Systemic therapy acts upon the entire organism including the bone marrow niche and our studies in p16-3MR aged mice demonstrate that genetic clearance of senescent cells can also improve BM MSC hematopoietic supporting function (Fig. 2). To assess whether short-term senolytic treatment with 753b impacts the bone marrow microenvironment, we performed single-cell RNA-sequencing on CD45⁻Lin⁻Ter119⁻ bone marrow stromal cells isolated from young, aged vehicle-treated, and aged 753b-treated mice (SFig. 7A). Unsupervised clustering identified a diverse set of stromal populations, including adipogenic and osteogenic mesenchymal stromal cell (MSC) subsets, fibroblast and myofibroblast populations, sinusoidal and arteriolar endothelial cells, osteoblasts, pericytes, and megakaryocytes (SFig. 7A–C). Cell-cycle inference analysis showed similar distributions of cycle phases across clusters in all groups, indicating no major changes in proliferation related to aging or short-term 753b treatment (SFig. 7D). Expression of *Bcl2* and/or *Bcl2l1* was strongly elevated in aged adipogenic-MSC_1 and chondrocytes, consistent with accumulation of senescent cells in the bone marrow niche with aging (SFig. 7E), which was reversed by 753b treatment, indicating effective senolysis (SFig. 7E). Comparative analysis of cluster proportions revealed a mild reduction of adipogenic MSCs and increase of adipocyte progenitor populations in aged_Veh mice relative to young controls. Short-term 753b treatment produced only modest shifts in stromal composition, generally preserving the stromal landscape (SFig. 7F). Taken together, these data indicate that the lineage composition and cycling are largely maintained in the bone marrow niche during physiological aging. Treatment with 753b achieves targeted senolysis without perturbing niche architecture.

**SFigure 7.**
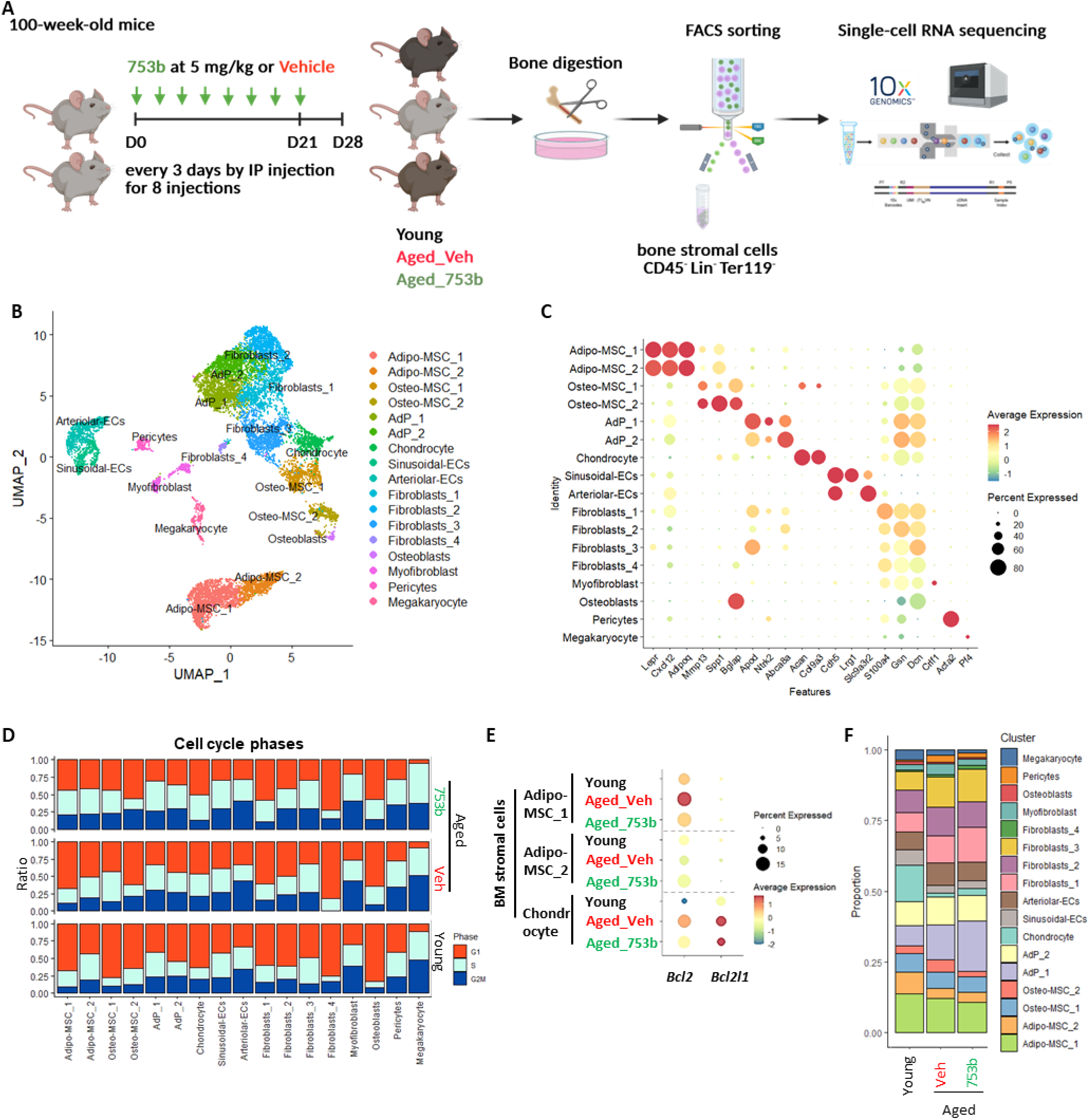
Short-term 753b treatment has little impact on bone marrow stromal cell composition and cycling. (A) Schematic of experimental design. (B) UMAP visualization of identified stromal cell populations, composite of all conditions. (C) Expression of representative marker genes used to annotate stromal clusters; dot size indicates the percentage of cells expressing the marker and color reflects average expression. (D) Distribution of cell-cycle phases (G1, S, G2/M) within each stromal cluster across groups. (E) Expression of *Bcl2* and *Bcl2l1* (*Bcl-xl*) in the three groups. (F) Relative proportions of each stromal cell type identified in the BM niche by scRNA-seq.

### Senescent cell clearance restores bone marrow cell-cell communication signaling networks altered in aging

A major function of the bone marrow niche is to send and receive critical paracrine cues to the hematopoietic compartment necessary to support continued blood production. To investigate age-related changes to cell-cell communication networks and their response to senescent cell clearance, we integrated our single-cell RNA-seq stromal and hematopoietic compartment profiles from young, aged vehicle-treated, and aged 753b-treated mice and projected them as joint UMAP profiles (Fig. 6A). Cell-cell communication networks were inferred using a reference dataset of 3,379 ligand-receptor pairs (CellChat V2^40^) and quantified by the total number of significant ligand-receptor interactions (edge count) and the overall communication probability (aggregated interaction strength) among all stromal and hematopoietic populations. We found a steep decline in communication connectivity between stromal and hematopoietic cell populations, with reduced number of significant interactions and lower overall signaling strength. 753b treatment partially restored both the complexity and strength of cell-cell interactions toward the young state (Fig. 6B). Pairwise analysis of ligand-receptor interactions between major cell types showed that aged bone marrow exhibited weakened connectivity, particularly within the stromal compartment and between major stromal and hematopoietic populations, while interactions within the hematopoietic compartment were preserved or even enhanced (Fig. 6C, left). 753b treatment partially reversed these changes (Fig. 6C, right, SFig. 8). In cluster-by-cluster CellChat analysis using young controls as a reference (Fig. 6D), aged bone marrow showed generalized attenuation of paracrine signals between niche cells (blue box) and from canonical niche populations (MSCs, chondrocytes, fibroblasts, and endothelial cells) to non-primed HSCs (pink box), accompanied by accentuated signaling from HSCs and myeloid-restricted progenitors (Fig. 6D, left; green and yellow boxes). 753b treatment restored many lost interactions originating from niche subsets and simultaneously relaxed tightened interactions within the aged hematopoietic compartments (Fig. 6D, middle), re-establishing a reciprocal communication landscape that more closely resembles the young BM signaling architecture (Fig. 6D, right).

Pathway-specific analysis identified aging-related disruption of key signaling routes, many of which are partially restored by senescent cell clearance. This included weakened extracellular matrix and adipokine-related interactions that are critical for niche architecture and metabolic support, such as laminin, adiponectin, and fibronectin-integrin (FN1) signaling (SFig. 9). Conversely, inflammation- and myeloid-related pathways such as Notch, App and Ifn-I signaling that were markedly enriched in aged BM could be substantially reversed by senolytic treatment (SFig. 10).

Chronic low-grade inflammation is a hallmark of tissue aging (termed “inflammaging”) that disrupts BM homeostasis through dysregulated cell-cell signaling networks. In the aged BM niche, senescent stromal cells secrete a variety of cytokines, chemokines, growth factors, and matrix-remodeling enzymes, known as senescence-associated secretory phenotype (SASP), that sustain chronic inflammatory stress and impair HSC maintenance. Hence, we next examined SASP-associated gene signatures in aged MSCs. Compared to young controls, aged MSCs displayed elevated expression of SASP-associated cytokines and chemokines (Fig. 6E), which was largely reversed by 753b treatment, indicating that senolysis can effectively suppress pathological MSC-derived inflammatory signaling and partially normalize the inflammatory milieu of the aged niche. Further pathway-level analysis revealed that IFN-I and TNF inflammatory signaling cascades were also highly enriched in aged BM (SFig. 11A). Expression of *Tnf* and *Ifn (*such as *Ifnb1, Ifna2*, and *Ifna16)* genes was highest in hematopoietic progenitors, particularly neutrophil precursors (Rep_2, pNeu_1 and pNeu_2) (SFig. 11B), indicating that hematopoietic cells themselves can act as dynamic components of the BM microenvironment and contribute substantially to intercellular communication. These inflammatory signals were markedly amplified with aging, especially interferon gene expression in neutrophil progenitors (Fig. 6F, SFig. 11C-D), whereas senolytic treatment attenuated inflammatory signatures, resembling a more young-like state.

Together, these results demonstrate that aging profoundly rewires BM cell-cell communication networks, weakening supportive stromal-HSC signaling interactions while amplifying inflammation. Hematopoietic cells, particularly neutrophil progenitors, emerge as key signaling hubs during aging beyond well-established niche components, providing an additional source of pro–inflammatory cues that reinforce BM inflammaging. Senolytic treatment with 753b normalizes aging-related changes in the cell-cell communication landscape by attenuating inflammatory processes in both niche and hematopoietic compartments, rebalancing and rejuvenating the global signaling ecosystem to support HSC function.

**Figure 6.**
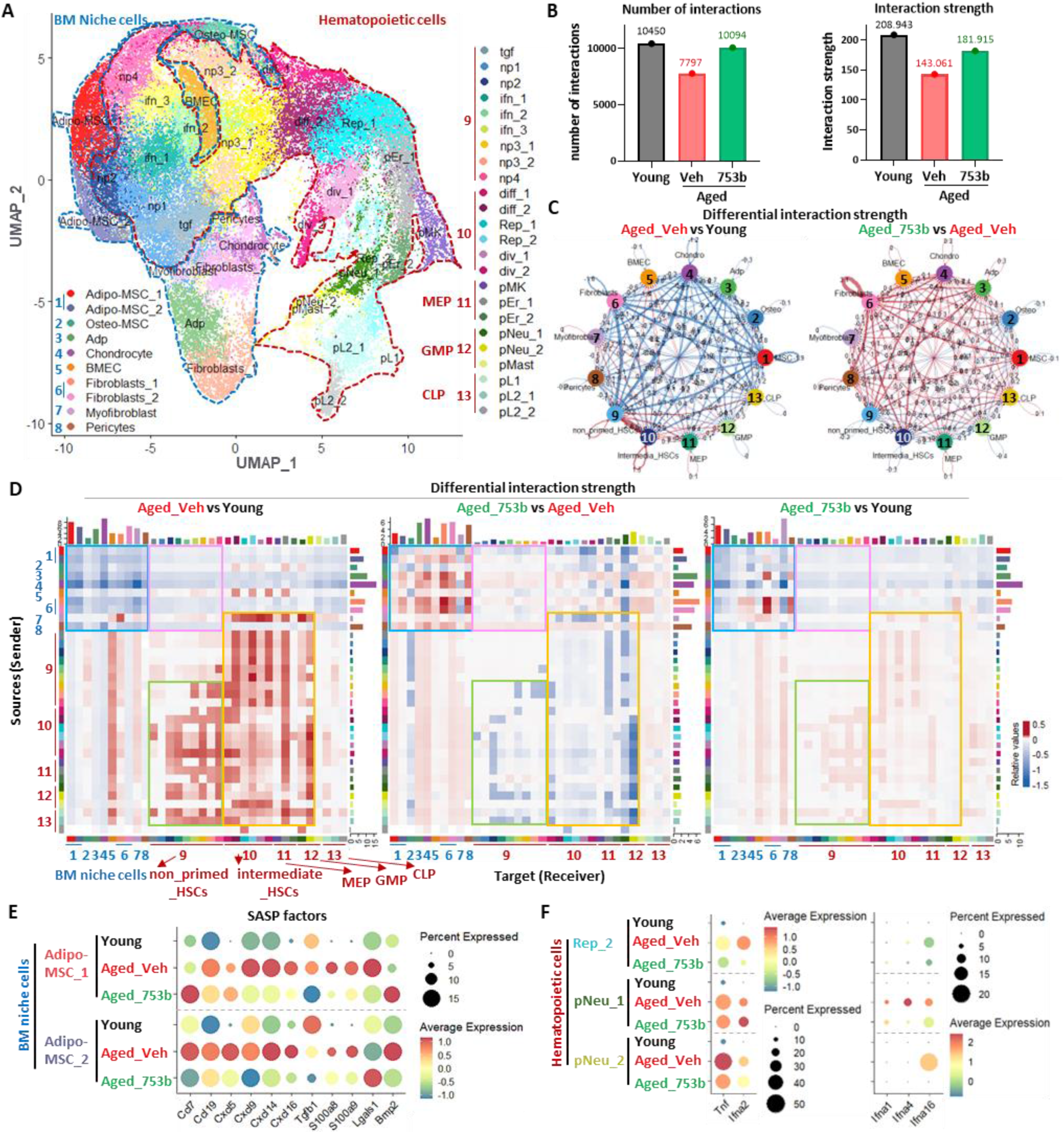
Senescent cell clearance restores BM intercellular signaling networks altered by aging. (A) UMAP visualization of integrated BM stromal and hematopoietic single–cell transcriptomes from young, aged vehicle–treated (Aged_Veh), and aged 753b–treated (Aged_753b) mice, with major BM niche populations (blue labels 1–8) and hematopoietic subsets (red labels 9–13) indicated. (1: mesenchymal stromal cells (MSCs); Adipo-MSC_1, Adipo-MSC_2. 2: Osteo-MSC. 3: Adipocyte progenitor; Adp. 4: Chondrocyte. 5: Bone marrow endothelial cells; BMEC. 6: Fibroblasts; Fibroblasts_1, Fibroblasts_2. 7: Myofibroblast. 8: Pericytes. 9: non_primed_HSCs; tgf, np1, np2, ifn_1, ifn_2, ifn_3, np3_1, np3_2, np4. 10: intermediate_HSCs; diff_1, diff_2, Rep_1, Rep_2, div_1, div_2. 11: MEP; pMK, pEr_1, pEr_2. 12: GMP; pNeu_1, pNeu_2, pMast. 13: CLP; pL1, pL2_1, pL2_2.) (B) Quantification of CellChat-inferred global communication networks showing the total number of significant ligand-receptor interactions (left) and the integrated interaction strength (overall communication probability, right) among all stromal and hematopoietic populations in each condition. (C) CellChat-derived aggregated communication networks comparing Aged_Veh vs Young (left) and Aged_753b vs Aged_Veh (right); edge width encodes total communication probability, with red edges indicating increased and blue edges indicating decreased interaction strength between conditions. (D) Differential interaction strengths between individual source (rows) and target (columns) clusters. Colored bars denote individual cell types, with each color corresponding to the cluster identities and palette shown in Fig. 6A. The bar height indicates the degree of change in interaction strength between the two conditions. (E) Expression levels of SASP–associated factors in Adipo–MSC subsets. (F) Expression levels of indicated genes in select hematopoietic subsets across conditions.

**SFigure 8.**
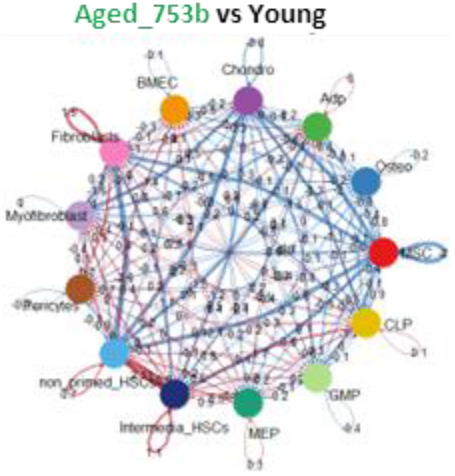
Clearance of senescent cell restores BM cell-to-cell signaling networks altered by aging. CellChat-derived aggregated communication networks comparing Aged_753b vs Young; edge width encodes total communication probability, with red edges indicating increased and blue edges indicating decreased interaction strengths between conditions.

**SFigure 9.**
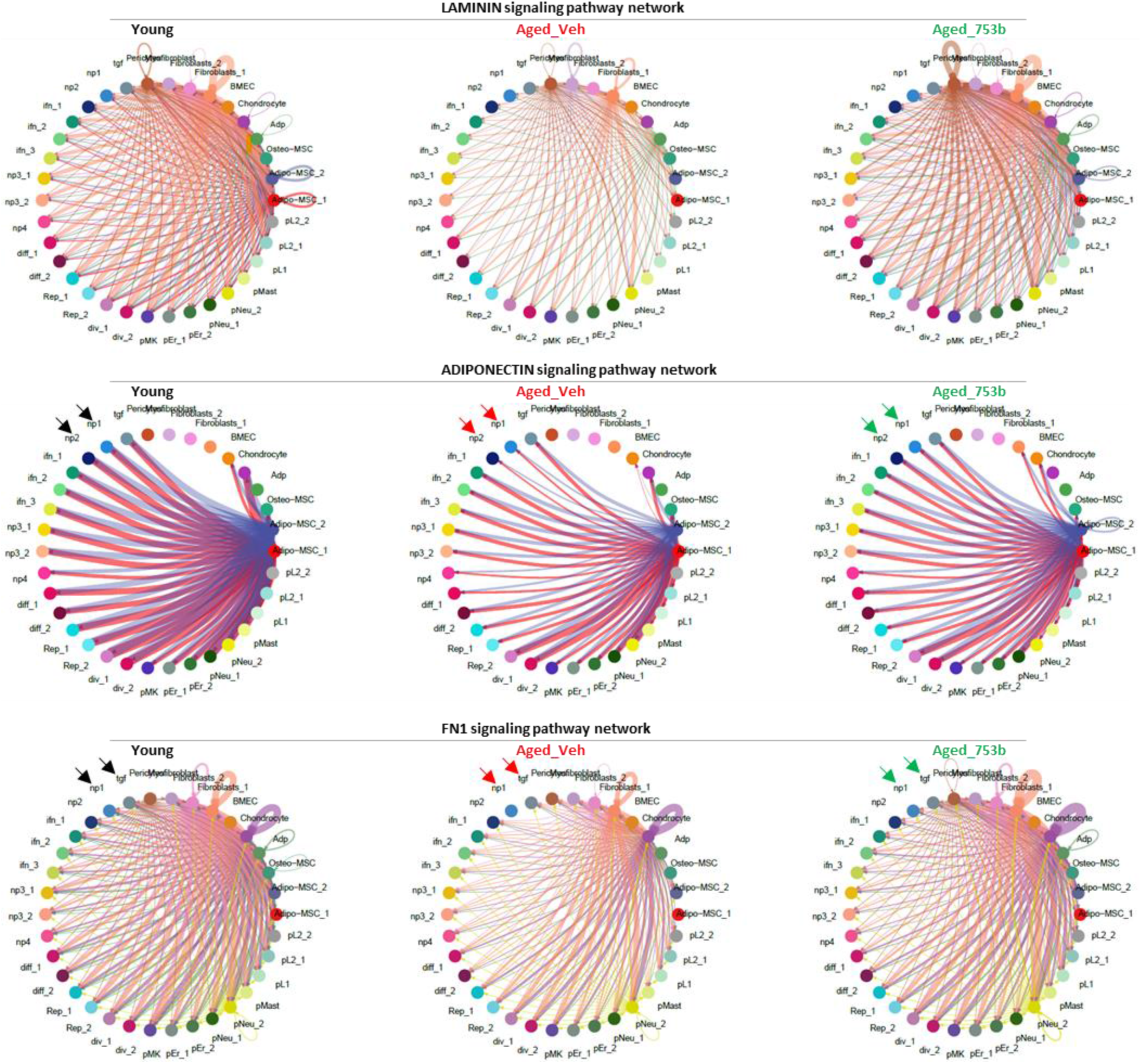
Signaling pathways that decline in aged bone marrow and are partially restored by senolytic treatment. Circle plots showing interaction strength among different cell populations across conditions. The edge colors are consistent with the sources as sender, and edge thickness is proportional to interaction strength, with thicker edges indicating stronger signaling.

**SFigure 10.**
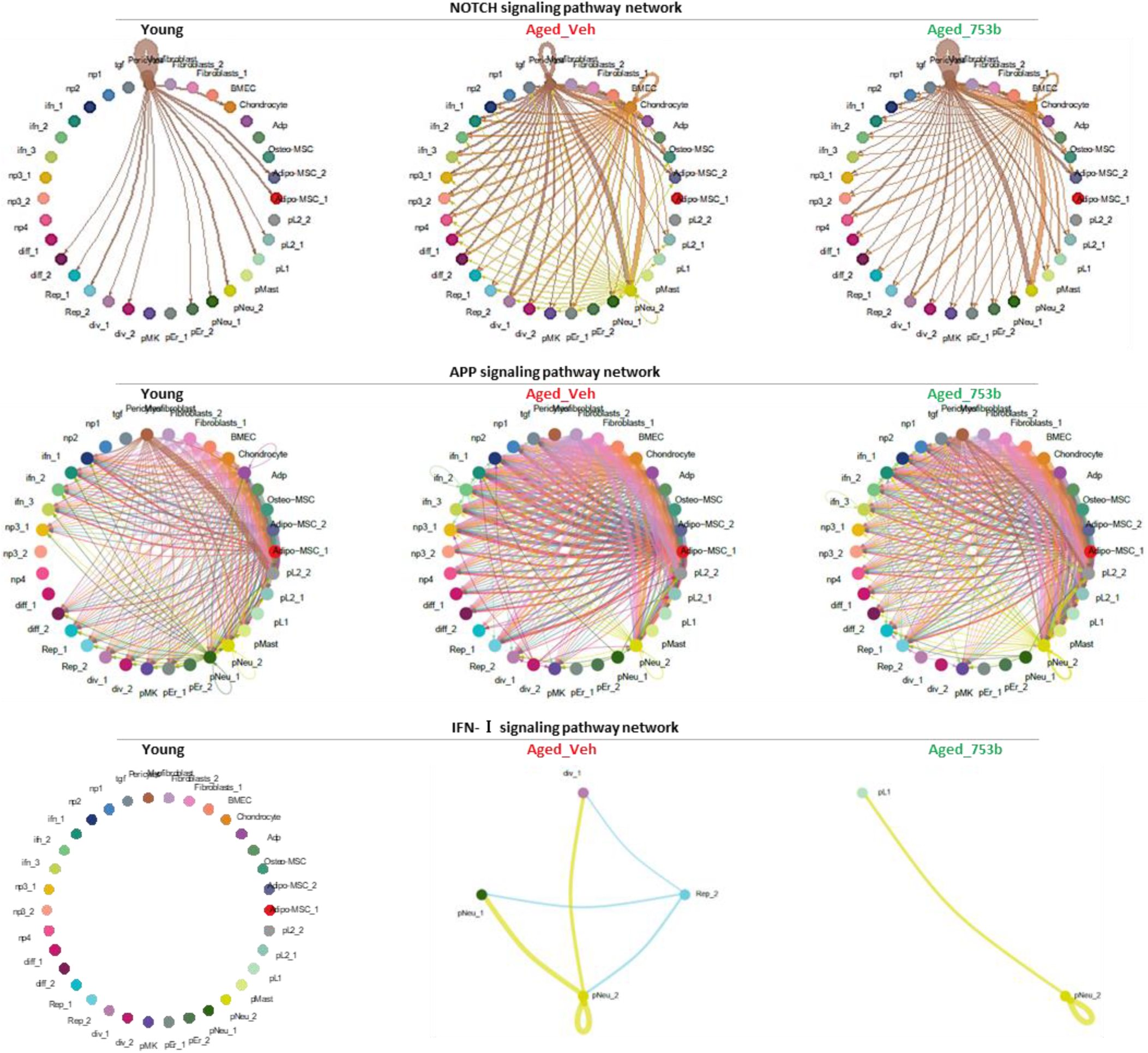
**Signaling pathways that elevated in aged bone marrow and are partially recovered by senolytic treatment.**

**SFigure 11.**
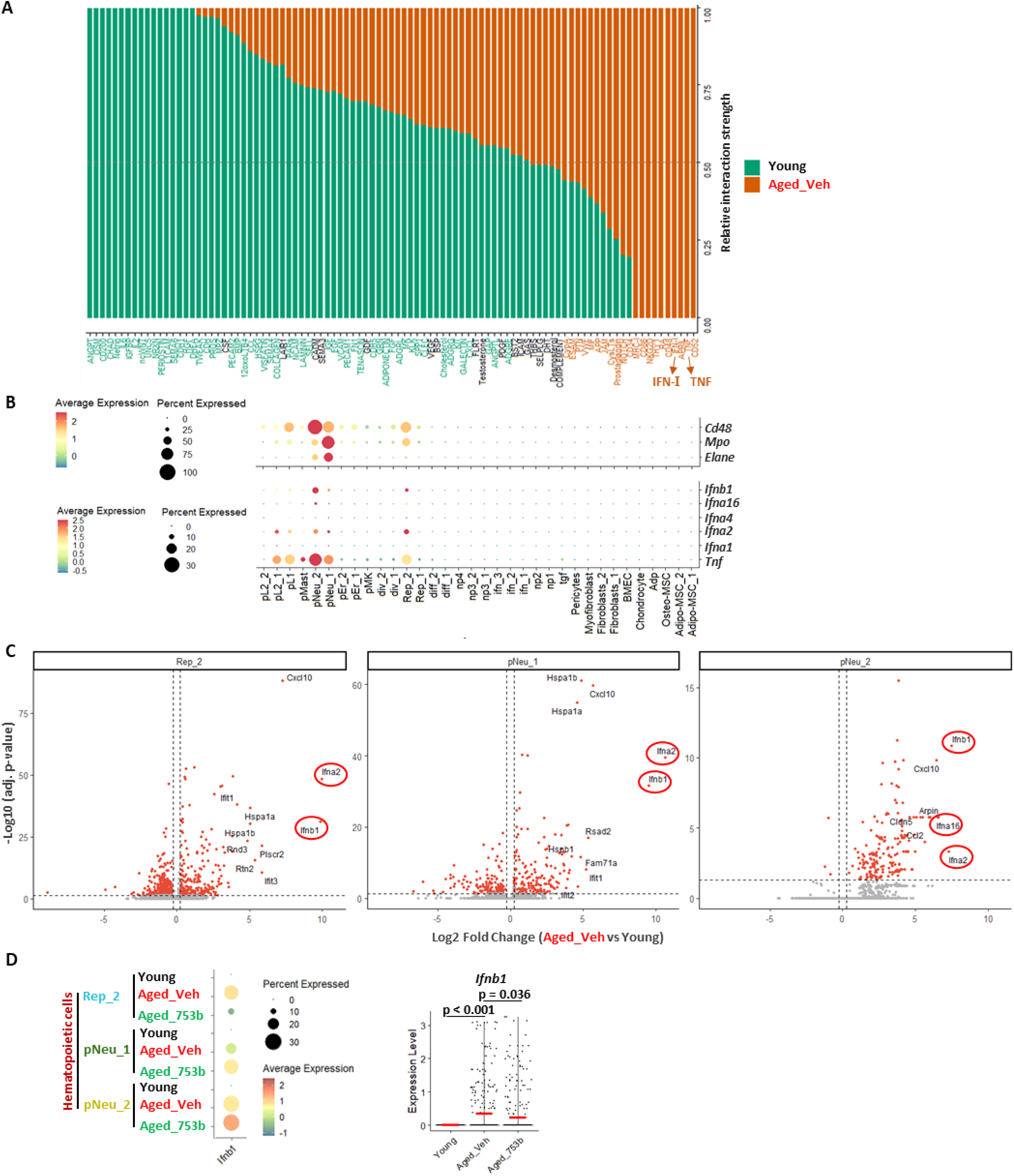
Elevated inflammatory signaling in aged hematopoietic cells. (A) Comparison of overall signaling pathway interaction strength between young and aged vehicle-treated groups. A paired Wilcoxon test was performed to assess significant differences in interaction strength between conditions. Pathways enriched in young mice are shown in green, and those enriched in aged vehicle-treated mice are shown in orange. (B) Expression levels of neutrophil linage and inflammatory genes across all clusters. (C) Volcano plots highlighting top elevated genes in indicated clusters in aged vehicle-treated group compared to young controls. (D) Integrated expression of the *Ifnb1* gene in Rep_2, pNeu_1 and pNeu_2 clusters (left) and transcript levels in individual cells of the 3 clusters (right) across conditions. Horizontal lines indicate medians; significance was determined using the Wilcoxon rank-sum test.

## Discussion

Aging in the hematopoietic system is accompanied by the accumulation of senescent cells, which contribute to a chronic pro-inflammatory milieu that progressively impairs hematopoietic function. This inflammatory burden likely disrupts stem and progenitor cell fitness, alters niche support and communications of BM MSCs and HSPCs, and accelerates functional decline across the compartment. Our study demonstrates that clearance of senescent cells in aged p16-3MR mice alleviates functional decline of HSCs and rejuvenates BM stromal cells, restoring hematopoietic support. These observations highlight the therapeutic promise of senolytic strategies, but also underscore the need for more selective and effective senolytics that can eliminate pathogenic senescent cells while preserving normal tissue integrity. Attributing to its characteristics as a highly effective next generation senolytic agent with improved hematologic safety profile, 753b has emerged as an attractive therapeutic candidate. Our study shows pharmacologic clearance of senescent cells with short-term 753b treatment reverses key molecular and functional hallmarks of hematopoietic aging. Single-cell RNA-seq data revealed that aged HSCs exhibited pronounced upregulation of stress and inflammatory pathways, including interferon and oxidative stress responses, alongside elevated aging-related gene expression signature. Derivation of an aging score from these transcriptional programs quantitatively captured the extent of HSC aging, showing a marked shift toward high-score populations in aged BM. Importantly, 753b significantly reduced these high-score states across multiple non-primed HSC subsets, transcriptionally rejuvenating the aged HSCs.

Consistent with molecular rejuvenation, 753b corrected functional imbalances in lineage differentiation potential that typify aged hematopoiesis. Aged HSCs showed increased expression of myeloid-biased genes and suppression of lymphoid specification factors, reflecting the classical myeloid skewing of aging. 753b treatment corrected age-associated myeloid gene overexpression while re-engaging lymphoid differentiation signatures, restoring HSC multipotency. Furthermore, upon transplant, BM from 753b-treated aged mice generated more balanced tri-lineage output than vehicle-treated aged controls.

In the BM niche, single-cell profiling of the stromal populations showed that global cellular composition and proliferative status were largely maintained during aging after short-term 753b treatment. Clearance of senescent cells by 753b attenuated SASP-mediated inflammatory signaling, restoring disrupted stromal-hematopoietic signaling networks and re-establishing extracellular matrix and adipokine-related interactions that support HSC maintenance. These data suggest that stromal rejuvenation synergizes with HSC-intrinsic remodeling to drive systemic restoration of hematopoietic aging.

Chronic IFN and TNF–α exposure accelerates HSC cycling, myeloid priming, and loss of lymphoid potential, ultimately leading to hematopoietic functional decline^41,42^. This study highlighted that hematopoietic cells themselves act as key signaling hubs, especially aged neutrophil progenitors exhibiting enhanced TNF and type I interferon signaling, providing an additional pro-inflammatory input that amplifies inflammaging. Senescent cell clearance by 753b markedly attenuated such self-reinforcing inflammatory circuit that perpetuate HSC aging.

Our analysis further highlights the importance of viewing BM function as an integrated system, in which dynamic communication among hematopoietic and niche cells underpins aging and rejuvenation processes. By targeting senescent cells, 753b not only reprograms intrinsic HSC states but also remodels the BM microenvironment, suggesting a feedback loop in which clearance of senescent cells in the niche promotes stem cell function and vice versa. Future studies incorporating higher-sensitivity multi-omic approaches will be helpful for delineating the precise molecular crosstalk between aged BM microenvironment and HSC dysfunction, ultimately allowing for more targeted therapeutic interventions.

The expanding aging population worldwide underscores the pressing need for clinic-appropriate rejuvenation agents that can preserve tissue fitness, mitigate age-related functional decline, and ultimately reduce frailty while extending healthspan. Current senolytic strategies have established proof of principle, but each approach remains constrained by important liabilities^43,44^. The first-generation, broadly acting agents dasatinib plus quercetin^20^ and fisetin^45^ offer the convenience of oral administration and clinical accessibility, yet their senolytic activity is comparatively modest, highly context dependent, with incomplete target coverage across senescent cell states, and toxicity. Navitoclax is mechanistically more potent through inhibition of BCL-2 family anti-apoptotic proteins^24^, but its clinical utility has been limited by on-target platelet toxicity, resulting in severe thrombocytopenia^28^, which constrains both dose intensity and treatment duration. More advanced cellular approaches such as senolytic CAR T-cells^46–48^ may improve specificity in principle, but they remain technologically complex, expensive, and difficult to deploy across heterogeneous senescent cell populations, where antigen selection and tissue access remain major barriers^44,49^. In addition, age-related immunosenescence may reduce CAR T cell treatment effectiveness in elderly patients^49^. Against this backdrop, 753b offers a distinct advantage by combining mechanistic potency with low toxicity. As a dual BCL-xL/BCL-2 PROTAC, 753b induces target degradation and promotes efficient clearance of senescent cells while avoiding the platelet toxicity of navitoclax. Notably, 753b reduced senescence burden, fibrosis, and MASH-driven hepatocarcinogenesis *in vivo*^30^, providing proof of concept for the efficacy and selectivity of senescent-cell elimination in alleviating chronic tissue injury, while also supporting an encouraging safety profile in a preclinical setting. In this sense, 753b represents a more translationally viable senolytic option than current pharmacologic or cell-based alternatives.

This study represents the first evaluation of 753b in aged animals in the context of BM function, where it exhibited a favorable safety profile and robust senolytic activity. Notably, its dual impact on both HSCs and stromal components led to improved hematopoietic function by attenuating age-associated myeloid skewing and inflammaging, two major contributors to age-related diseases and hematologic malignancies. These findings position 753b as a promising candidate for restoring hematopoietic balance without compromising stem cell integrity or niche architecture.

It is also important to note that while short-term 753b treatment produced meaningful molecular and functional rescue, the use of 100-week-old animals may have missed the optimal window for mitigating aging-induced HSC defects. While some level of correction is observed, it remains far from complete recovery. Previous reports show the hallmarks of aging in HSCs and BM microenvironment emerge at middle age^2,9,50^. Therefore, future studies exploring the impact of initiating senolytic interventions in mid-life, before irreversible stem cell exhaustion or niche remodeling occurs, and extending the treatment duration, could provide more insights into the temporal window and dosing strategies required for optimal rejuvenation of hematopoietic function.

Together, these results place senescent cell burden as a central, targetable determinant of hematopoietic and niche aging. By integrating senolytic intervention with BM single-cell analysis, this study provides direct evidence that clearance of senescent cells can recalibrate both intrinsic HSC transcriptional states and extrinsic niche communication. The ability of 753b to restore HSC proliferation, repopulation capacity, and lineage balance underscores the therapeutic potential of senolytic agents for rejuvenating aged BM function. Future studies optimizing the timing of senolytic initiation, dosing, treatment duration, and combinations with regenerative stimuli will be critical for translating 753b into clinical strategies for hematopoietic rejuvenation and mitigation of age–related disease.

## Methods

### Mice

All animal experiments were conducted in accordance with ethical guidelines and were approved by the Institutional Animal Care and Use Committee (IACUC) at the University of Arkansas, University of Florida (UF) and Johns Hopkins University. All recipient mice used for bone marrow transplantation in this study were C57BL/6.SJL (CD45.1, JAX stock # 002014) immune-competent, healthy females procured from the Jackson Laboratory.

### p16-3MR transgenic mice

p16-3MR transgenic mice were kindly provided by Drs. Judith Campisi and Marco Demaria, and maintained as previously described^13,17^. These mice carry a 3MR (trimodality reporter) fusion protein under the control of the *p16* (also referred to *Cdkn2a*, cyclin-dependent kinase inhibitor 2a) promoter^13^. The 3MR transgene encodes a fusion protein consisting of Renilla luciferase (LUC), monomeric red fluorescent protein (mRFP), and truncated herpes simplex virus 1 (HSV-1) thymidine kinase (HSV-TK), which converts ganciclovir (GCV) into a toxic DNA chain terminator to selectively kill HSV-TK-expressing senescent cells. To clear senescent cells in p16-3MR mice, 12-month-old male p16-3MR mice randomly divided in two groups (6 mice per group) and treated with vehicle or GCV (G2536, Sigma, USA) diluted in phosphate-buffered saline (PBS). GCV was administered by intraperitoneal (i.p.) injection at 25 mg/kg (0.1 ml) per day for 5 consecutive days per cycle, with two cycles separated by a 2-week interval till they reached 24 months of age. A group of untreated young (2-month of age) male C57BL/6J mice (JAX stock # 000664) was included as control. One week after the last treatment, mice were euthanized by CO2 suffocation followed by cervical dislocation. Inguinal fat tissues were harvested for RNA extraction to analyze expression of *Cdkn2* and several SASP factors mRNA as described later.

### 753b treatment

C57BL/6 mice of different ages (young, 2 months; naturally aged, 22 months) were obtained from the National Institute on Aging and group housed with ad libitum access to food and water. For 753b treatment, naturally aged mice were randomly assigned to receive either vehicle (10% ethanol, 30% PEG-400, 60% PHOSAL 50 PG; n = 6-8) or 753b (5 mg/kg in the same vehicle) by intraperitoneal injection every 3 days for a total of eight doses, as outlined in Fig. 3A. A cohort of untreated young mice served as controls. Seven days after the final injection, mice were euthanized by CO₂ asphyxiation followed by cervical dislocation, and tissues were collected for analysis.

### Isolation of bone marrow cells

Bone marrow cells were isolated from femurs and tibias. Briefly, mice were euthanized, hind limbs were dissected, and surrounding muscle and connective tissue were removed. Both ends of each bone were cut, and bone marrow was flushed with ice-cold PBS containing 2% FBS. The suspension was passed through a 70 µm cell strainer, centrifuged, and red blood cells were removed by ammonium-chloride-based lysis, followed by washing and resuspension in staining buffer (PBS, 2% FBS) for downstream flow cytometry (Aria II cell sorter, BD Biosciences, USA), FACS (Lin^-^, CD45R/B220^-^ CD3ε^-^ CD11b^-^Gr-1^-^ Ter-119^-^; HSCs, CD150^+^CD48^-^Lin^-^Sca1^+^c-Kit^+^; LT-HSCs, CD34^-^CD48^-^CD150^+^Lin^-^Sca1^+^c-Kit^+^).

### Analysis of p16 and phosphorylated p38 (p-p38) staining in HSC

Lin^−^ cells were stained with antibodies against HSC cell-surface markers and then fixed and permeabilized using Fixation-Permeabilization Solution (BD-Pharmingen, San Diego, CA). Subsequently, the cells were stained with anti-p16 (F–12, Santa Cruz Biotechnology, USA) and goat anti-mouse IgG-Alexa Fluor 488 (Thermo Fisher Scientific, USA), anti-p-p38-Alexa Fluor 647 (28B10, Cell Signaling Technology, USA), and 7-aminoactinomycin D (7-AAD) and then analyzed by flow cytometry as previously described^17^.

### Analysis of phosphor-histone H2AX (γ-H2AX) staining in HSC

Freshly sorted HSCs (∼2,000 cells per sample) were cytocentrifuged onto glass slides for immunostaining. Cells were fixed with 4% paraformaldehyde (PFA) for 15 min at room temperature and permeabilized in 0.1% Triton X–100 for 30 min. After PBS washes and blocking in PBS containing 1% bovine serum albumin (BSA; Sigma, USA) for 60 min, cells were incubated overnight at 4°C with primary antibodies against γ–H2AX-FITC (2F3, Biolegend, USA). Following washing, appropriate fluorophore-conjugated secondary antibodies were applied, and nuclei were counterstained with DAPI (Sigma, USA). Images were acquired using an Axioplan research microscope (Carl Zeiss Inc., Germany). For each slide, >100 cells were scored across >30 random fields to quantify γ-H2AX foci for calculating the average number of γ-H2AX foci per cell.

### Competitive repopulation assay

Fifty LT-HSCs were isolated from young, vehicle-treated old p16-3MR, and GCV-treated old p16-3MR mice, and mixed with 3 × 10^5^ competitive bone marrow cells pooled from three young CD45.1 mice. The cell mixture was then transplanted into lethally irradiated (9.5 Gy) CD45.1 recipients via retro-orbital injection of the venous sinus. Donor-cell engraftment in the recipients was analyzed at 4 months after transplantation as previously described^17^.

### Isolation of bone mesenchymal stromal cells (MSCs)

As described previously^17^, Femur, tibia and pelvic bones were harvested from euthanized mice, cleaned of surrounding soft tissue, and flushed to remove bone marrow cells. The bones were then cut into small (<1-mm) fragments and digested in PBS containing 15% FBS, 0.1% collagenase I, and 15 μg/ml DNase I (Sigma) at 37 °C for 1 h with gentle agitation. The released cells were collected, bone fragments were removed, and the suspension was filtered through a 40-μm cell strainer. CD45^−^Lin^−^PI^−^ bone stromal cells were collected by cell sorting.

### Co-culture of LSK cells with MSCs

Isolated MSCs were seeded into wells of 12-well plates at 7.5×10^4^/well with 2 ml cobblestone area-forming cell (CAFC) assay medium (10% heat inactivated fetal bovine serum, 5% heat inactivated hours serum, 5 x 10^-5^ M 2-mercapital-ethanol, 0.48 mg/ml hydrocortisone, 100 U/mL penicillin, and 100 μg/mL streptomycin). After 24h culture, the wells were refreshed with 2 ml fresh CAFC medium after removing old medium and dead cells. Five thousand LSK cells from young, vehicle-treated old p16-3MR, and GCV-treated old p16-3MR mice were sorted into each well as illustrated in Figure 2B. Half of the cell culture medium in each well was replaced with fresh CAFC medium after 3-day co-culture. Three days later, both suspension cells and adherent cells were collected from each well of the co-culture and resuspended into 2 ml MethoCult™ GF M3434 medium (StemCell Technologies, Canada) and 0.5 ml of the cells in MethoCult™ GF M3434 medium was seeded into wells of 12-well plates using a 1 ml tuberculin syringe fitted with a 16-gauge blunt needle. Numbers of granulocyte-macrophage progenitor cells (CFU-GM) and multi-potential granulocyte, erythroid, macrophage, and megakaryocyte progenitor cells (CFU-GEMM) were counted on day 7 and 12 culture, respectively, and expressed as number of CFU-GM and CFU-GEMM per 10^3^ input LSK cells from each group.

### Single cell RNA sequencing (scRNA-seq)

Single cells were barcoded using the 10× Genomics Chromium single-cell platform, and cDNA libraries were prepared following the manufacturer’s instructions (Chromium Single Cell 3’ Kits v3.1; 10× Genomics, USA). Hematopoietic stem cells were isolated via FACS as Lin^-^c-Kit^+^Sca1^+^CD150^+^ cells, while stromal cells were collected from the CD45^-^Lin^-^Ter119^-^ populations from both male and female animals (n ≥ 6 per group). Live cell counts were determined using a hemocytometer. The prepared cells were subsequently loaded onto a 10x Genomics Chip, aiming for an output of approximately 10,000 cells per sample. The pooled libraries were then sequenced using the NovaSeq 6000 S4 system (Illumina, USA), targeting ∼400 million reads per library. Count matrices generation was conducted using the 10x Genomics Cell Ranger pipeline (version 5.0.0), following the 10x Genomics guidelines. The demultiplexed FASTQ files were aligned to the mm10 reference genome.

Downstream analysis, including data normalization, integration, and clustering was conducted using the Seurat package (version 4.3.0)^37^. Low-quality cells were excluded based on the following criteria: fewer than 500 detected genes, fewer than 500 unique molecular identifiers (UMIs), or >5% mitochondrial transcript content. Cell clusters were manually annotated and refined according to established marker gene expression profiles reported in prior studies^33,51–53^. Communication probabilities and ligand-receptor interactions were inferred and visualized using the CellChat R package (version 2.2.0)^40^.

The scRNA-seq dataset has been deposited to GEO (accession number GSE316278).

## Statistics

All statistical analyses were performed with GraphPad Prism version 10.3.1. Unpaired comparisons between two groups were performed using Student’s *t*-test for normally distributed data with equal variances, Welch’s correction for unequal variances. Statistical significance of gene expression levels in violin plots for scRNA-seq data was assessed using the Wilcoxon rank-sum test. *p* values less than 0.05 were considered statistically significant.

## Acknowledgements

This work was supported by the National Institutes of Health grants R01AG063801 (DZ, GZ), R01CA211963 and P30AG013319 (DZ), and R01DK121831 (OAG), Ocala Royal Dames for Cancer Research, Inc. (BY), Dr. Nihal Tumer Junior Faculty Research Catalyst Award (BY), and the American Cancer Society (ACS) Institutional Research Grant to the University of Florida Health Cancer Institute (UFHCI). UFHCI is an NCI-designated cancer center (P30CA247796). Flow cytometry, gene expression and genotyping, next-generation sequencing, and bioinformatics analyses were performed at the University of Florida Interdisciplinary Center for Biotechnology Research (ICBR) Cytometry Core (CY, RRID:SCR_019119), Gene Expression & Genotyping Core (GE, RRID:SCR_019145), NextGen DNA Sequencing Core (NS, RRID:SCR_019152), and Bioinformatics Core (BI, RRID:SCR_019120).

## Statements and Declarations

DZ, GZ and PZ are inventors on patent applications for use of BCL-xL/2 PROTACs as senolytic agents. DZ and GZ are co-founders of and have equity in Dialectic Therapeutics, which develops BCL-xL PROTACs to treat cancer. All other authors have nothing to declare.

Study was conceptualized by Daohong Zhou, Bowen Yan, Olga Guryanova, Jennifer Elisseeff, and Ying Liang. The animal model, treatment, husbandry, tissue distribution and material preparation were performed by Bowen Yan, Han Jin, Yang Yang, Jianhui Chang, Ha-Neui Kim, Maria Almeida, Prabhjot Kaur, and Qingchen Yuan. 753b was synthesized and provided by Peiyi Zhang and Guangrong Zheng. Data collection and analysis were performed by Bowen Yan, Ying Liang, and Daohong Zhou. Next generation sequencing data was analyzed by Jason Brant, Bowen Yan and Kalyanee Shirlekar. Marco Demaria provided p16-3MR mice and reviewed and edited the manuscript. The first draft of the manuscript was written by Bowen Yan and all authors commented on previous versions of the manuscript. The Funding was acquired by Daohong Zhou, Guangrong Zheng, Olga Guryanova, and Bowen Yan. All figures created using BioRender were produced under a full BioRender license that permits publication and distribution. All authors read and approved the final manuscript.

