## Supplemental File for "Targeting cellular senescence alleviates bone marrow aging"

**SUPPLEMENTARY INFORMATION**

**Supplementary Figure 1**

Clearance of senescent cells mitigates HSC functional decline in naturally aged p16-3MR mice

**Supplementary Figure 2**

753b targets senescent cells with high efficacy and low toxicity

**Supplementary Figure 3**

Clearance of senescent cells with 753b reverses myeloid expansion contributed by aged HSCs

**Supplementary Figure 4**

Clearance of senescent cells with 753b reshapes the transcriptomes of aged HSCs towards a more youthful composition of subpopulations

**Supplementary Figure 5**

Consolidating aging score in independent mouse and human HSCs datasets

**Supplementary Figure 6**

753b treatment ameliorates aging‑associated transcriptomic features in HSCs

**Supplementary Figure 7**

Short‑term 753b treatment has little impact on bone marrow stromal cell composition and cycling

**Supplementary Figure 8**

Clearance of senescent cell restores BM cell-to-cell signaling networks altered by aging

**Supplementary Figure 9**

Signaling pathways that decline in aged bone marrow and are partially restored by senolytic treatment

**Supplementary Figure 10**

Signaling pathways that elevated in aged bone marrow and are partially recovered by senolytic treatment

**Supplementary Figure 11**

Elevated inflammatory signaling in aged hematopoietic cells

Supplementary Methods

**Supplementary figures**

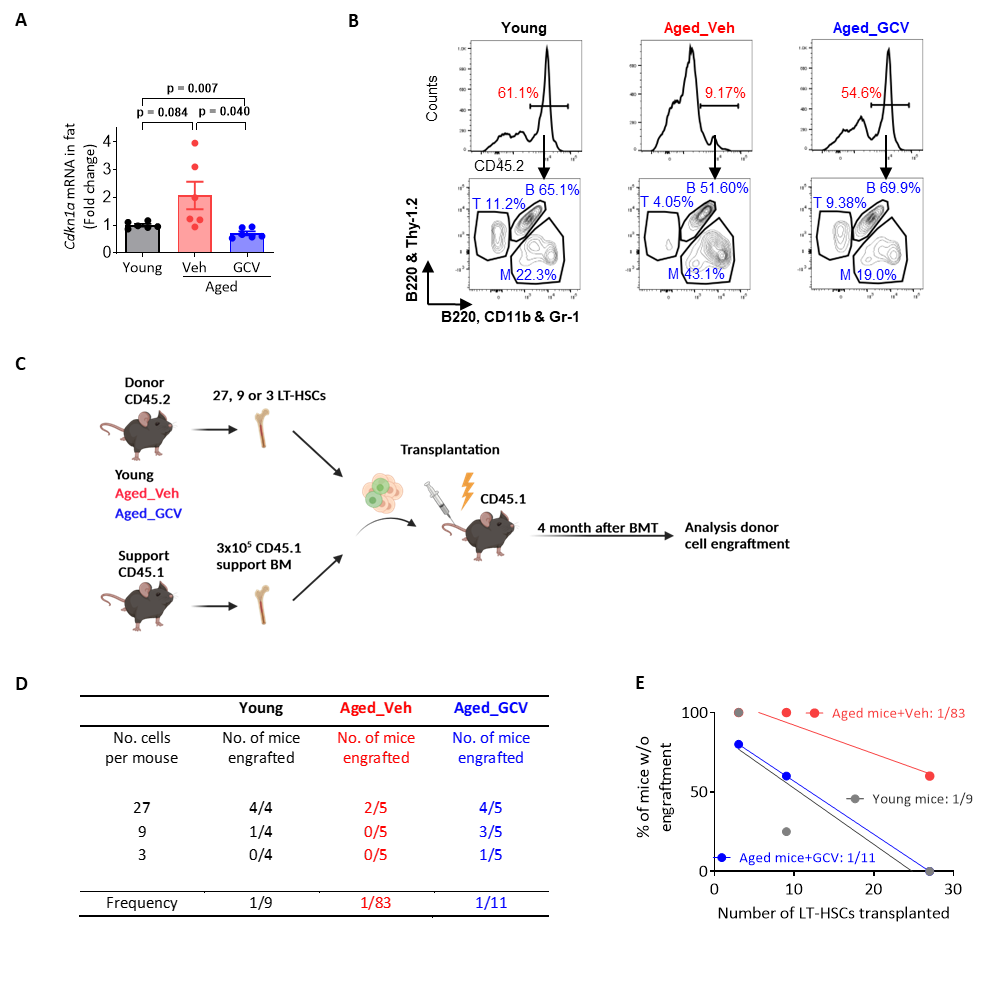

**SFigure 1. Clearance of senescent cells mitigates HSC functional decline in naturally aged p16-3MR mice.**

(A) *Cdkn1a* transcript levels in adipose tissue (qRT-PCR); mean ± SEM, Student’s *t*-test with Welch’s correction. (B) Flow cytometry testing donor chimerism (CD45.2%) and peripheral blood lineage composition (T cells, B cells, myeloid cells) at 4 months after BMT using donor material from young, aged vehicle-treated, or aged GCV-treated mice. (C-E) Design of the limiting dilution transplantation assay (C), frequency of reconstitution (D), and negative binomial regression analysis of the percentage of non-engrafted mice versus number of immunophenotypic LT-HSCs transplanted (E).

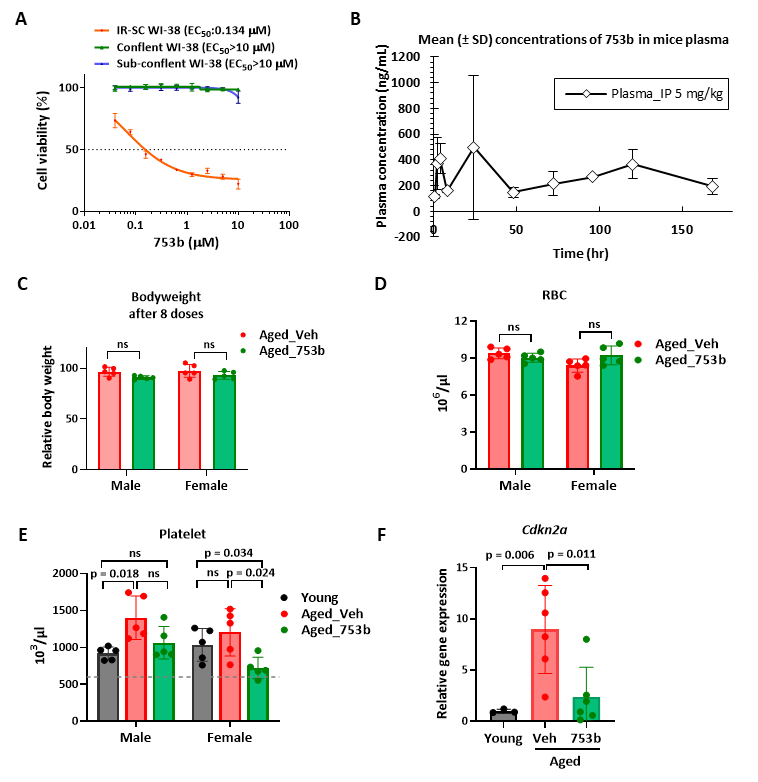

**SFigure 2. 753b targets senescent cells with high efficacy and low toxicity.**

(A) Dose-response curves to 753b in confluent, sub-confluent and senescent (IR-SC) WI-38 cells (IR-SC) after 72 hours of treatment; mean ± SD. (B) Plasma concentration of 753b in mice following a single IP injection (5 mg/kg). Plasma samples were measured at 0.5, 2, 4, 8, 24, 48, 72, 96, 120 and 168 hours post injection; mean ± SD. (C) Body weight of aged mice after 8 doses of 753b or vehicle in both sexes; mean ± SEM; Student’s *t*-test. (D) Red blood cell (RBC) counts in peripheral blood of aged male and female mice treated with vehicle or 753b; mean ± SEM; Student’s *t*-test. (E) Platelet counts in young, aged vehicle-treated, and aged 753b-treated mice; mean ± SEM; Student’s *t*-test. (F) Relative gene expression of *Cdkn2a* (coding for p16) in spleens of young, aged vehicle-treated, and aged 753b-treated mice; mean ± SEM; Student’s *t*-test with Welch’s correction.

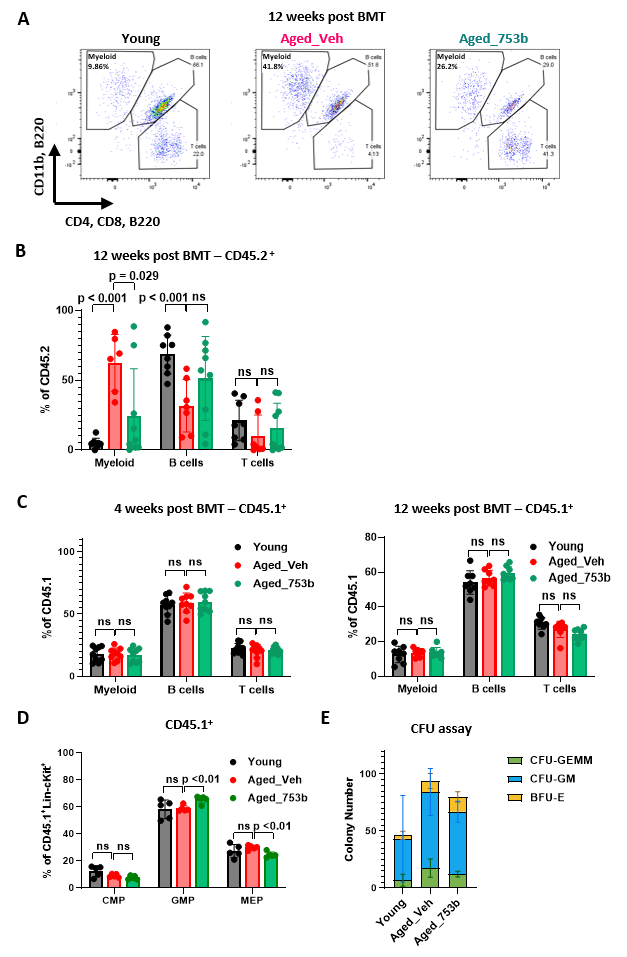

**SFigure 3. Clearance of senescent cells with 753b reverses myeloid expansion contributed by aged HSCs.**

(A, B) Representative flow cytometry plots of peripheral blood leukocytes 12 weeks after BMT (A) and quantification of myeloid, B, and T cells in CD45.2^+^ donor-derived compartment (B). (C) Myeloid, B, and T cell percentages in wild-type control competitor-derived CD45.1^+^ compartment 4 and 12 weeks post-BMT; mean ± SEM; Student’s *t*-test. (D) Frequencies of CMP, GMP and MEP populations in competitor-derived CD45.1^+^ compartment; mean ± SEM; Student’s *t*-test. (E) CFU assay showing the numbers of colony types across indicated groups; mean ± SD.

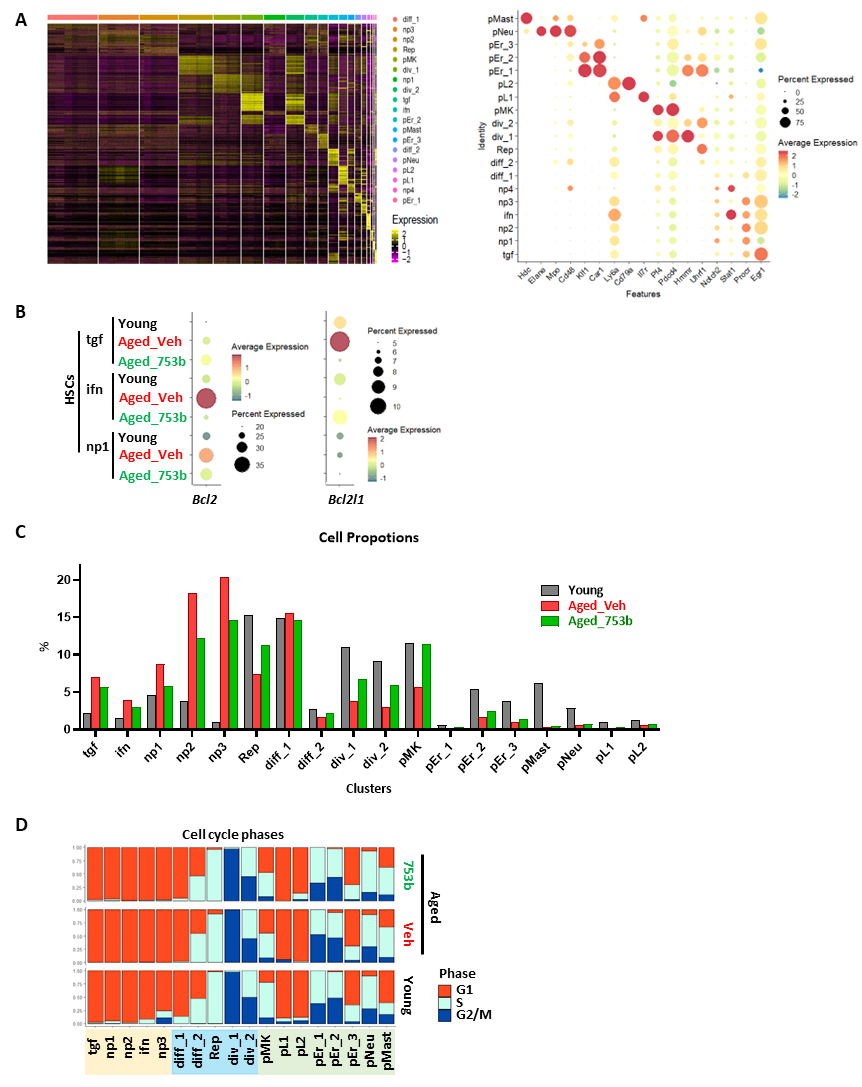

**SFigure 4. Clearance of senescent cells with 753b reshapes the transcriptomes of aged HSCs towards a more youthful composition of subpopulations.**

(A) Left, heatmap of scaled expression values for the top marker genes defining each cell cluster identified in single-cell RNA seq data. Right, dot plot showing representative cluster‑specific marker gene expression levels, where dot size represents the percentage of cells expressing each gene and color indicates average expression level. (B) *Bcl2* and *Bcl2l1* (*Bcl‑xL*) expression levels in tgf, ifn, and np1 HSC clusters across conditions. (C) Cluster proportions across conditions. (D) Distribution of cell cycle phases (G1, S, G2/M) across clusters and conditions.

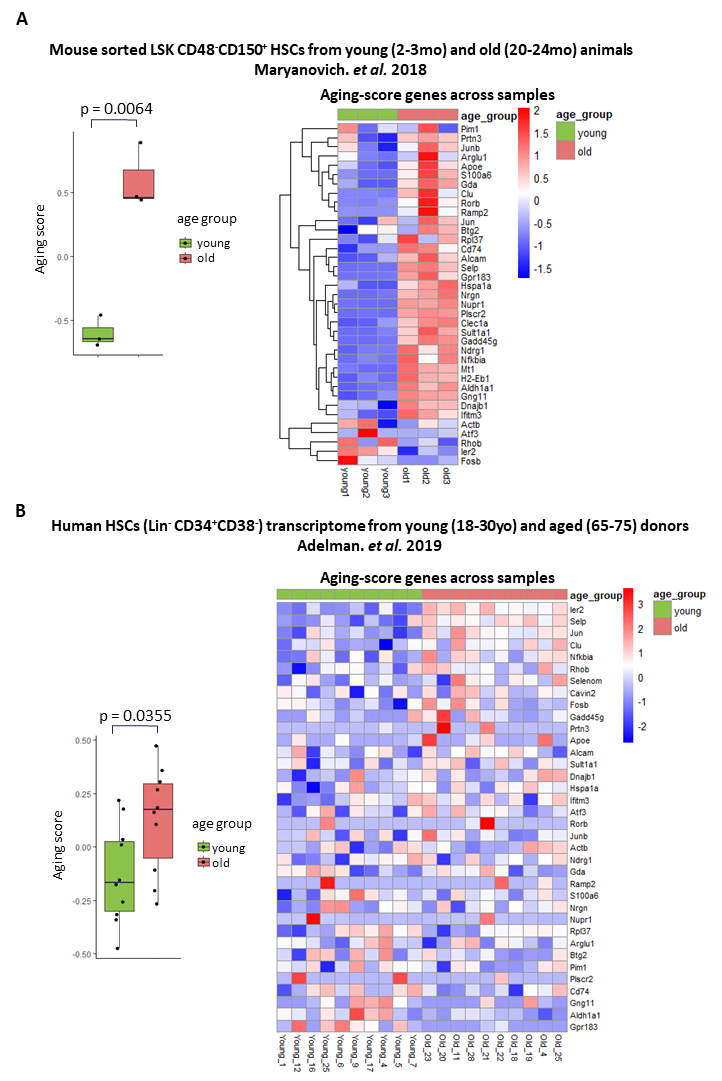

**SFigure 5. Consolidating aging score in independent mouse and human HSCs datasets.**

Boxplot (left) comparing age groups using the aging score derived from our single-cell–based aging gene set, calculated for published bulk transcriptomes of sorted murine LSK CD48^-^CD150^+^ HSCs from young (2-3mo) and old (20-24mo) mice (A) and human HSC (Lin^-^CD34^+^CD38^-^) transcriptomes from young (18-30yo) and aged (65-75) donors (B). Heatmap (right) shows z‑scaled expression of aging-score genes across samples with annotation bars indicating age groups.

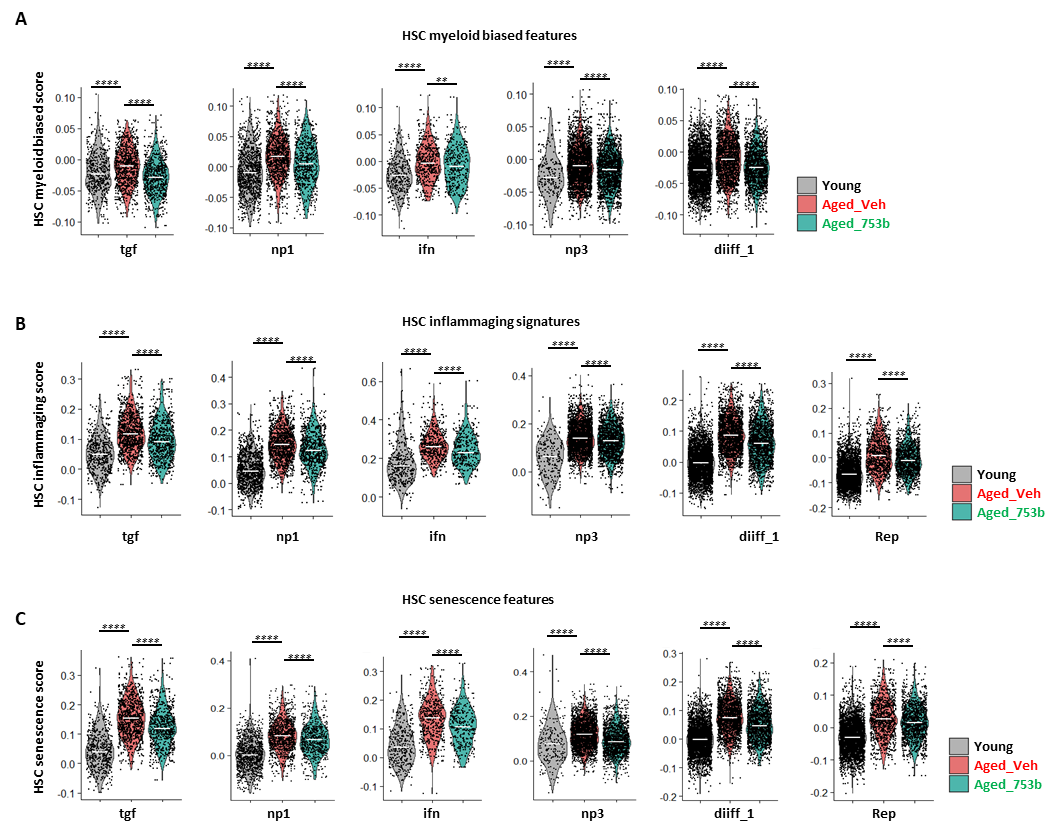
**SFigure 6. 753b treatment ameliorates aging‑associated transcriptomic features in HSCs.**

(A–C) Gene signature scores for HSC myeloid-biased features (A), HSC inflammaging signatures (B), and HSC senescence features (C) across the indicated clusters in young, aged_veh and aged_753b mice. Each point represents a single cell; violin width reflects the distribution of scores within each group, and horizontal lines denote the median, Wilcoxon rank-sum test.

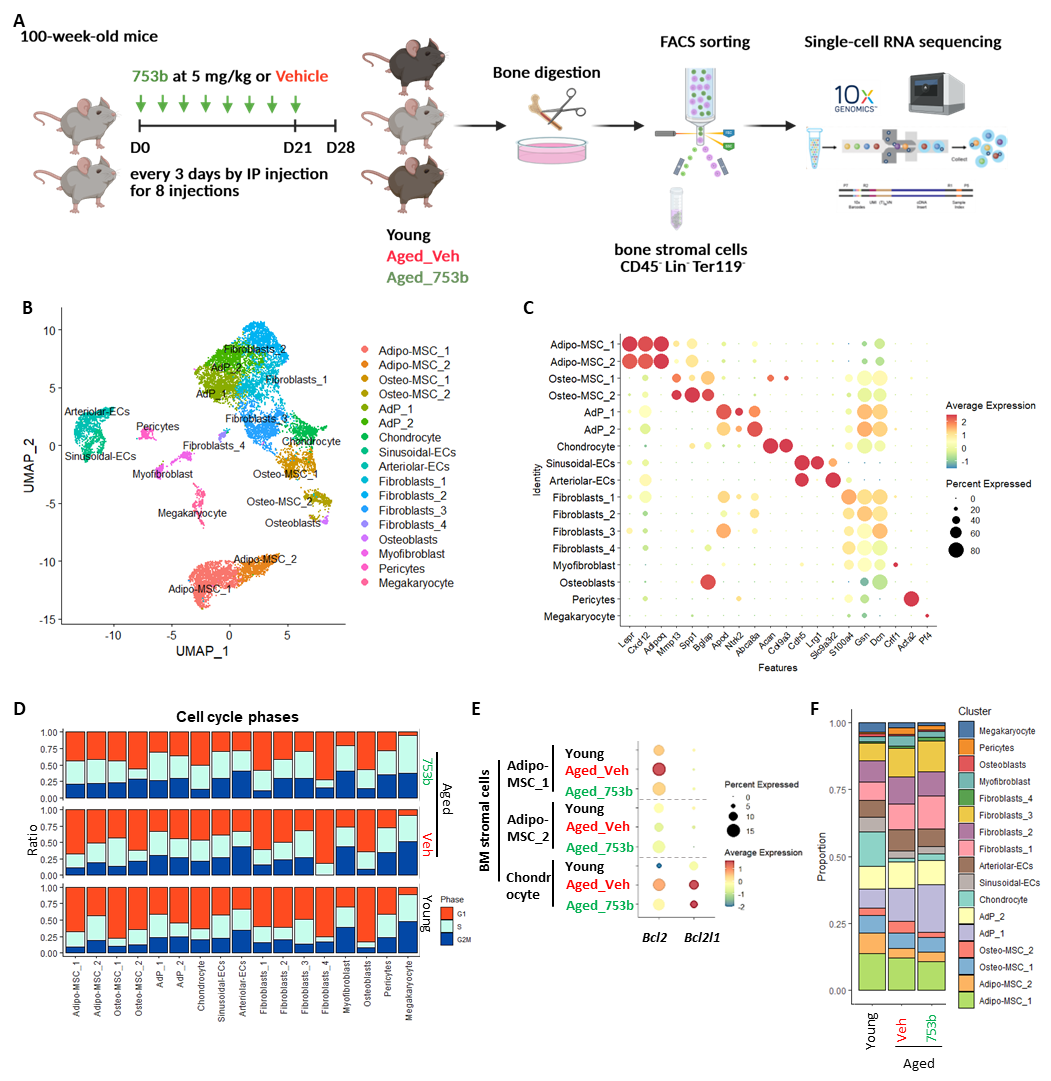

**SFigure 7. Short‑term 753b treatment has little impact on bone marrow stromal cell composition and cycling.**

(A) Schematic of experimental design. (B) UMAP visualization of identified stromal cell populations, composite of all conditions. (C) Expression of representative marker genes used to annotate stromal clusters; dot size indicates the percentage of cells expressing the marker and color reflects average expression. (D) Distribution of cell‑cycle phases (G1, S, G2/M) within each stromal cluster across groups. (E) Expression of *Bcl2* and *Bcl2l1* (*Bcl‑xl*) in the three groups. (F) Relative proportions of each stromal cell type identified in the BM niche by scRNA-seq.

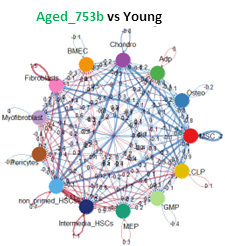

**SFigure 8. Clearance of senescent cell restores BM cell-to-cell signaling networks altered by aging.**

CellChat‑derived aggregated communication networks comparing Aged_753b vs Young; edge width encodes total communication probability, with red edges indicating increased and blue edges indicating decreased interaction strengths between conditions.

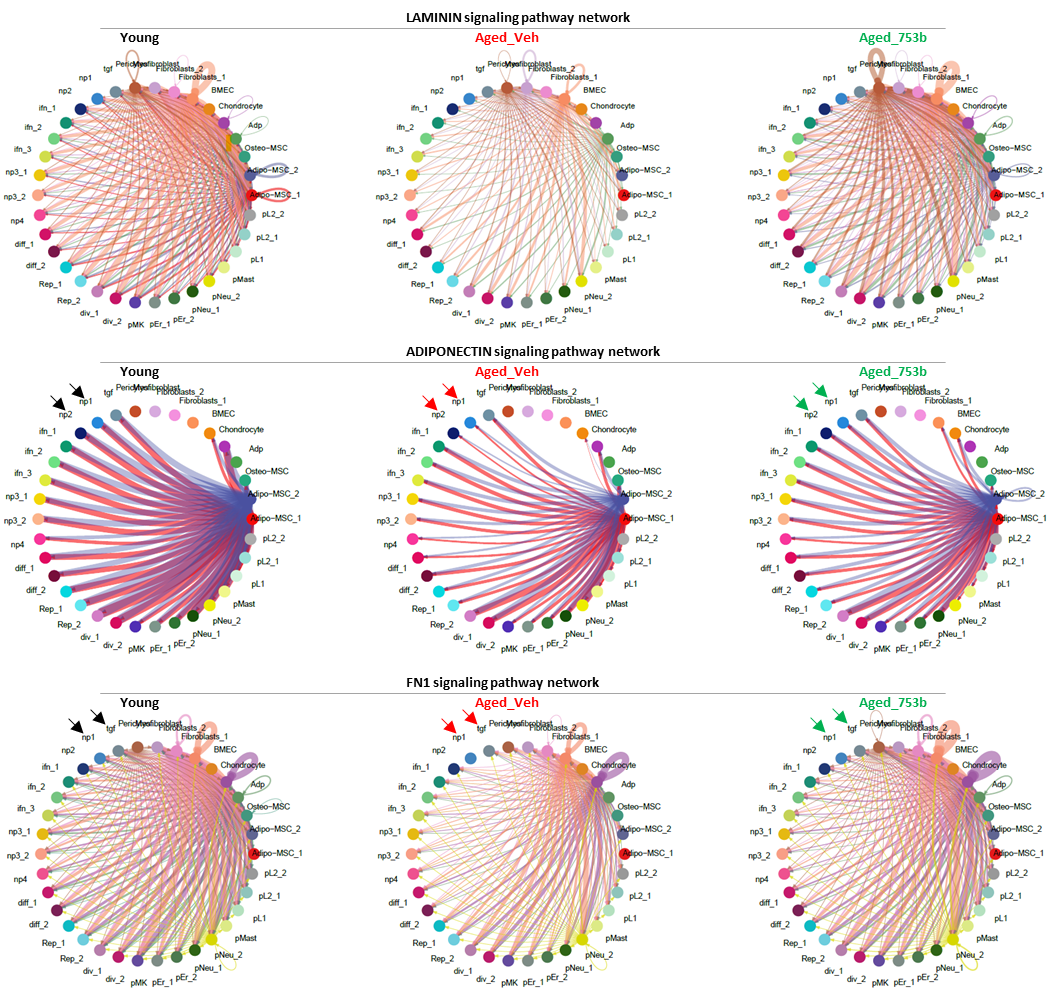

**SFigure 9. Signaling pathways that decline in aged bone marrow and are partially restored by senolytic treatment.**

Circle plots showing interaction strength among different cell populations across conditions. The edge colors are consistent with the sources as sender, and edge thickness is proportional to interaction strength, with thicker edges indicating stronger signaling.

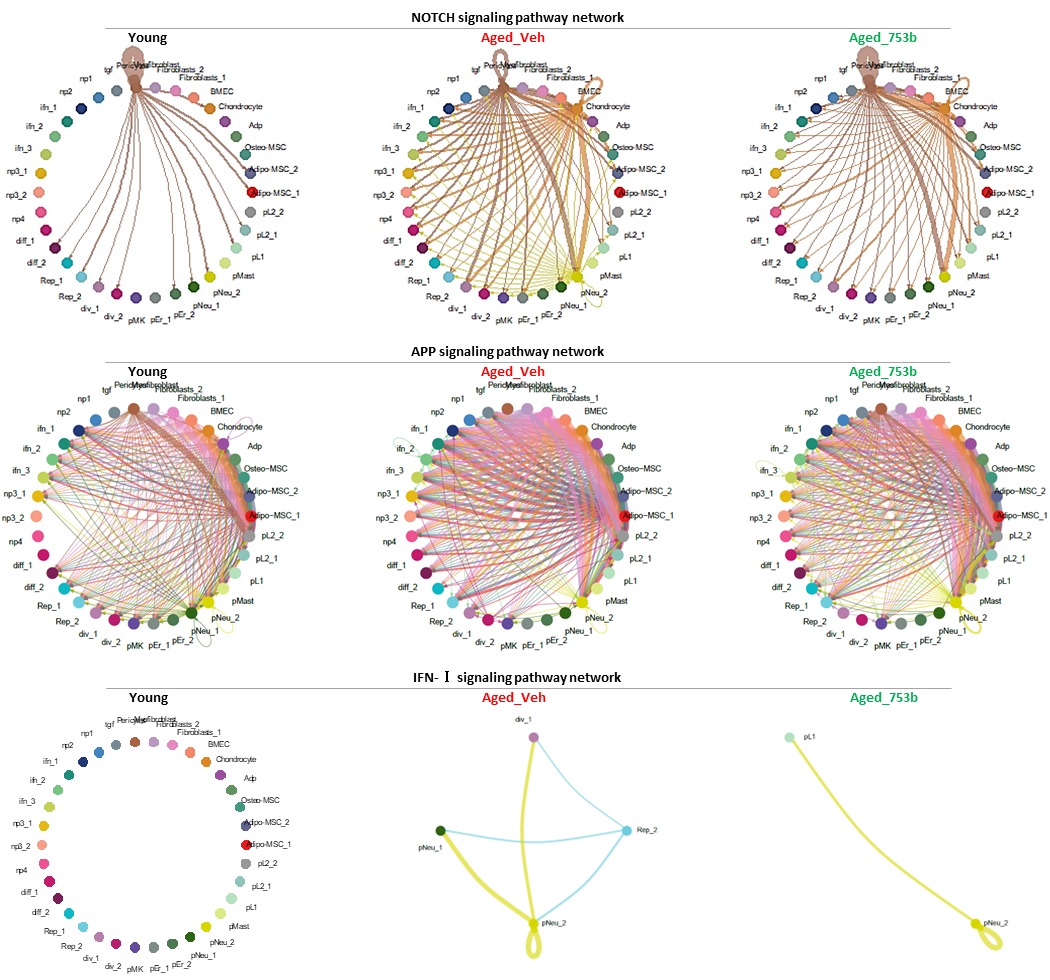

**SFigure 10. Signaling pathways that elevated in aged bone marrow and are partially recovered by senolytic treatment.**

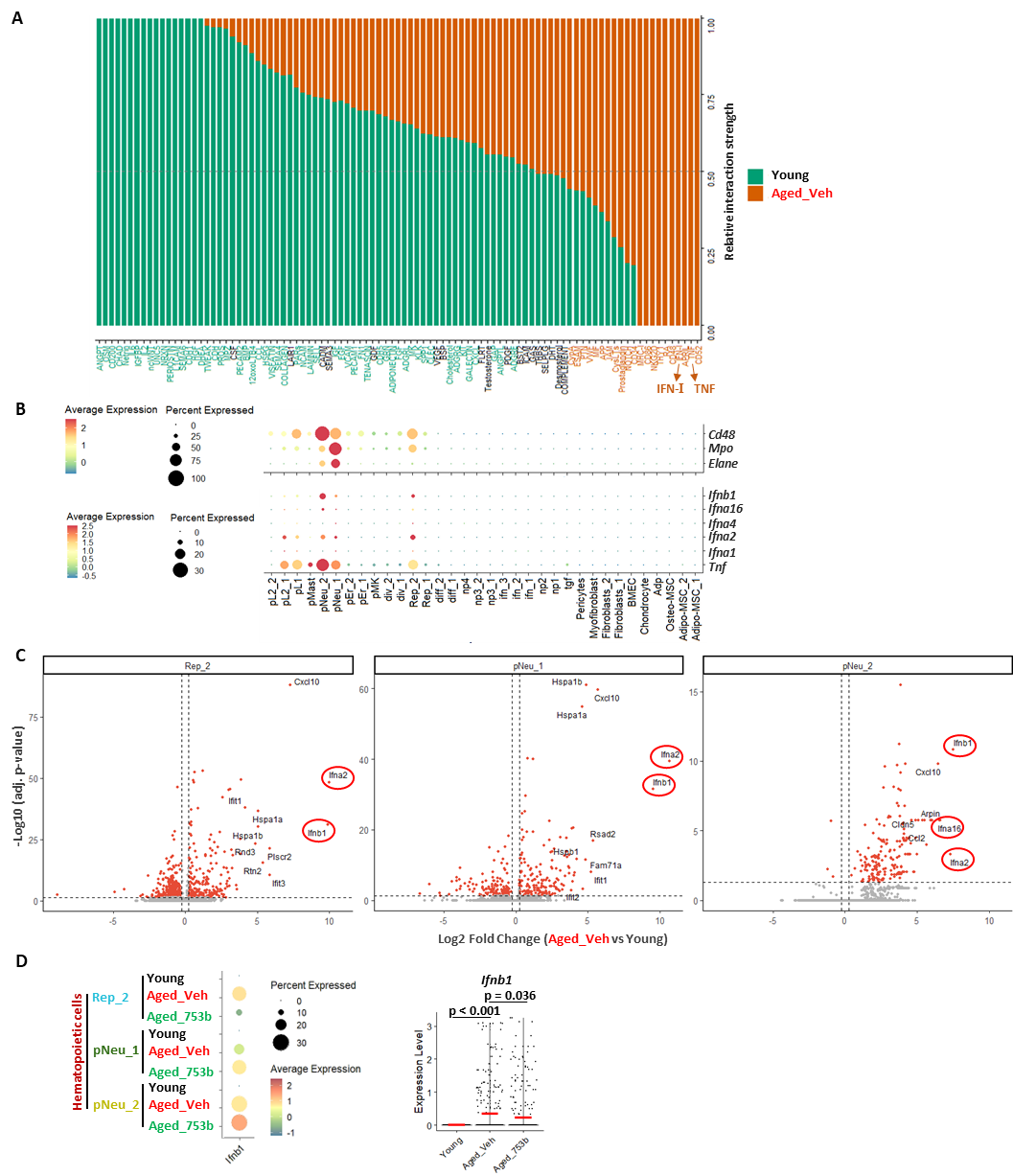

**SFigure 11. Elevated inflammatory signaling in aged hematopoietic cells.**

(A) Comparison of overall signaling pathway interaction strength between young and aged vehicle-treated groups. A paired Wilcoxon test was performed to assess significant differences in interaction strength between conditions. Pathways enriched in young mice are shown in green, and those enriched in aged vehicle-treated mice are shown in orange. (B) Expression levels of neutrophil linage and inflammatory genes across all clusters. (C) Volcano plots highlighting top elevated genes in indicated clusters in aged vehicle-treated group compared to young controls. (D) Integrated expression of the *Ifnb1* gene in Rep_2, pNeu_1 and pNeu_2 clusters (left) and transcript levels in individual cells of the 3 clusters (right) across conditions. Horizontal lines indicate medians; significance was determined using the Wilcoxon rank-sum test.

**Supplementary Methods**

**Limiting dilution transplantation assay**

Long-term hematopoietic stem cells (LT-HSCs) were isolated from donor bone marrow by FACS as Lin^-^Sca-1^+^c-Kit^+^CD150^+^ cells. Specified cell numbers of LT-HSCs (3, 9, or 27 per recipient) were resuspended in PBS together with 3 × 10^5^ supporting whole bone marrow cells from congenic CD45.1 mice and transplanted into lethally irradiated (9.5 Gy) CD45.1 recipient mice via retro-orbital injection of the venous sinus.

Peripheral blood samples were collected at 16 weeks post-transplantation to assess donor chimerism and lineage output by flow cytometry. Mice were designated as successfully engrafted if ≥1% multilineage donor-derived (CD45.2^+^) cells were detected in peripheral blood across myeloid and lymphoid lineages. Engraftment frequencies at each input dose were analyzed using ELDA software or negative binomial regression to estimate functional HSC frequency.

**Analysis of blood counts**

Approximately 50 μl of whole blood was collected from mice via the submandibular plexus into EDTA-coated microtainer tubes (Sarstedt, Germany), and complete blood counts were measured using an automated hematology analyzer (HEMAVET, Drew Scientific, USA).

**Flow cytometry analysis**

The blood and bone marrow single-cell suspensions were lysed with ammonium‑chloride-based lysis buffer and then resuspended in PBS containing 0.2% BSA. The cells were then stained with fluorochrome-conjugated antibodies and analyzed by multiparameter flow cytometry performed using LSR Fortessa instrument (BD Biosciences, USA). Data were analyzed using FlowJo software (v10.4.1). The following antibodies were used:

| **Antibody** | **Fluorophore** | **Company** | **Catalog number** | **Dilution** |
| --- | --- | --- | --- | --- |
| CD45R / B220 | purified | BD Biosciences | 553084 | 1:200 |
| CD3ε | purified | BD Biosciences | 550275 | 1:200 |
| CD11b | purified | BD Biosciences | 557394 | 1:200 |
| Gr-1 | purified | BD Biosciences | 550291 | 1:200 |
| Ter-119 | purified | BD Biosciences | 550565 | 1:200 |
| CD45R / B220 | biotin | BD Biosciences | 553086 | 1:200 |
| CD3e | biotin | BD Biosciences | 553060 | 1:200 |
| CD11b | biotin | BD Biosciences | 553309 | 1:200 |
| Gr-1 | biotin | BD Biosciences | 553125 | 1:200 |
| Ter-119 | biotin | BD Biosciences | 553672 | 1:200 |
| CD16 / CD32 | Purified | BD Biosciences | 553142 | 1:200 |
| CD45.2 | FITC | BD Biosciences | 553772 | 1:100 |
| CD45R / B220 | APC | BD Biosciences | 561880 | 1:200 |
| CD45R / B220 | PE | BD Biosciences | 553089 | 1:200 |
| CD11b | PE | BD Biosciences | 557397 | 1:200 |
| Gr-1 | PE | BD Biosciences | 553128 | 1:200 |
| Streptavidin | FITC | Biolegend | 405201 | 1:200 |
| Sca1 | PE | BD Biosciences | 553336 | 1:100 |
| Sca1 | PE-Cy™ 7 | BD Biosciences | 558162 | 1:100 |
| c-kit | APC-H7 | BD Biosciences | 560250 | 1:100 |
| c-kit | APC-eFluor® 780 | eBioscience | 47-1171-82 | 1:100 |
| CD150 | APC | eBioscience | 17-1501-81 | 1:100 |
| CD34 | Alexa Fluor® 700 | eBioscience | 56-0341-82 | 1:20 |
| CD48 | Pacific blue | Biolegend | 103418 | 1:200 |
| CD45 | APC | BD Biosciences | 559864 | 1:200 |
| CD11b | APC | BD Biosciences | 553312 | 1:200 |
| Sca-1 | Alexa Fluor® 700 | eBioscience | 56-5981-82 | 1:200 |
| CD45.1 | PacBlue | BioLegend | 110722 | 1:200 |
| CD45.2 | APC | BioLegend | 109814 | 1:200 |
| CD4 | FITC | BioLegend | 100406 | 1:200 |
| CD8 | FITC | BioLegend | 100706 | 1:200 |
| B220 | FITC | BioLegend | 103206 | 1:200 |
| B220 | APC/Cyanine7 | BioLegend | 103224 | 1:300 |
| Ter-119 | APC/Cyanine7 | BioLegend | 116223 | 1:300 |
| CD3 | APC/Cyanine7 | BioLegend | 101222 | 1:300 |
| CD4 | APC/Cyanine7 | BioLegend | 100414 | 1:300 |
| CD11b | APC/Cyanine7 | BioLegend | 101226 | 1:300 |
| Gr-1 | APC/Cyanine7 | BioLegend | 108424 | 1:300 |
| CD19 | APC/Cyanine7 | BioLegend | 115530 | 1:300 |
| NK1.1 | APC/Cyanine7 | BioLegend | 108724 | 1:300 |
| c-Kit | APC | BioLegend | 105812 | 1:200 |
| Sca1 | BV605 | Biolegend | 108134 | 1:200 |
| CD150 | PE | BioLegend | 115904 | 1:200 |
| CD16/CD32 | AlexaFluor700 | eBioscience | 48-0161-82 | 1:200 |
| CD34 | FITC | eBioscience | 11-D341-85 | 1:200 |
| CD45.1 | PE-Cy7 | BioLegend | 110730 | 1:200 |
| CD45.2 | PerCP-Cy5.5 | BioLegend | 109828 | 1:200 |
| CD48 | PacBlue | BioLegend | 103418 | 1:200 |
| CD45 | APC/Cyanine7 | eBioscience | 47-0451-82 | 1:200 |
| CD31 | APC | eBioscience | 17-0311-82 | 1:200 |
| Sca1 | PE/Cyanine7 | BD Biosciences | 558162 | 1:100 |
| CD51 | PE | eBioscience | 12-0512-83 | 1:100 |

**753b Pharmacokinetic (PK) study**

The PK study was performed by BioDuro Inc. (Irvine, USA). Male C57BL/6 mice (7-9 weeks of age) were given 753b (5 mg/kg in 60 % Phosal 50 PG, 30% PEG-400 and 10% ethanol) via intraperitoneal (IP) injection. Blood was collected from the mice at 0.5, 2, 4, 8, 24, 48, 72, 96, 120 and 168 hours post a single injection and analyzed for plasma concentrations of 753b using a pre-validated mass spectrometry method.

**Quantitative real-time PCR (RT-PCR)**

Total cellular RNA was extracted from cells or approximately 25 mg frozen mouse tissue using RNeasy Mini kits (74106, QIAGEN, Germany), and reverse transcription was performed as described previously^1,2^. All reactions were performed in triplicate on an ABI StepOnePlus Real-Time PCR System (Applied Biosystems, USA). The primers used in this study are listed as follows.

Sequences of the primers used for qRT-PCR

| Gene | Forward sequences | Reverse sequences |
| --- | --- | --- |
| Mouse *Cdkn2a* | CGGTCGTACCCCGATTCAG | GCACCGTAGTTGAGCAGAAGAG |
| Mouse *Il1a* | CCATAACCCATGATCTGGAAGAGAC | GTCCACATCCTGATATATAGTTTG |
| Mouse *Il1β* | CCATAACCCATGATCTGGAAGAGAC | GTCCACATCCTGATATATAGTTTG |
| Mouse *Il6* | CTGCAAGAGACTTCCATCCAG | AGTGGTATAGACAGGTCTGTTGG |
| Mouse *Tnfa* | TGAACTTCGGGGTGATCGGTC | CACTTGGTGGTTTGCTACGACG |
| Mouse *Ccl5* | CCCGCACCTGCCTCACCATATGG | CCTTCGAGTGACAAACACGACTG |
| Mouse *Hprt* | AGCAGTACAGCCCCAAAATGGTTA | TCAAGGGCATATCCAACAACAAAC |
